# IL-17A primes an early progenitor compartment to tune the erythropoietic feedback circuit

**DOI:** 10.1101/2024.12.16.628761

**Authors:** Qiu C Wu, Aishwarya Swaminathan, Ashley Winward, Logan Lalonde, Yung Hwang, Noah Littman, Merav Socolovsky, Allon M Klein

## Abstract

Feedback control of erythropoiesis exemplifies conflicting goals in tissue homeostasis: maintaining fast reactivity to stress while minimizing proliferative burden on progenitors in the steady state. Here we show that these conflicting goals are tuned through the combinatorial action of cytokines. We find that IL-17A, a pro-inflammatory cytokine, mediates striking synergism with the negative feedback signal erythropoietin (Epo) in vivo, accelerating the erythropoietic response to hypoxia. A model of erythropoietic control shows increased reactivity may occur through two cell circuit designs, with one having far lower constitutive progenitor burden in normoxia. IL-17A acts through this optimal design by sensitizing progenitors to Epo, a model supported by multiple experimental observations. We suggest that IL-17A signals impending hypoxia during infections, tuning erythropoiesis in favor of a faster stress response. Our study highlights IL-17A as a potential erythropoietic therapeutic agent and serves as a model of homeostatic tuning in stem and progenitor cell circuits.

## Introduction

Homeostatic control of renewing adult tissues is maintained through negative feedback: a change in cell number or function elicits a signaling response that opposes and reverses the change. This control principle is simple, but the feedbacks themselves can be complex. Defining the cell circuits underlying feedback control provides a mechanistic understanding of regenerative biology. It has practical implications for controlling progenitor self-renewal and differentiation and for understanding their failure in aging and disease.

A property of many feedback control systems is that their performance can be optimized by tuning - a process whereby feedback circuit parameters are adjusted in anticipation of changing future demand (**Fig. 1A**). The goals of tuning may be to maximize the speed or accuracy with which systems sense and react to their environment, or to minimize resting-state fluctuations or ongoing costs in energy and resources^1-3^. In an optimized system, improving one objective may be at the expense of another (a Pareto Front, **Fig. 1B**)^4,5^. Well studied examples include changes to body temperature, blood glucose, blood calcium^6-8^. Tissue stem and progenitor cells might also tune performance to prioritize different performance objectives, and understanding how they do so could help to elucidate the complexity of cell circuits in tissue renewal.

**Figure 1:**
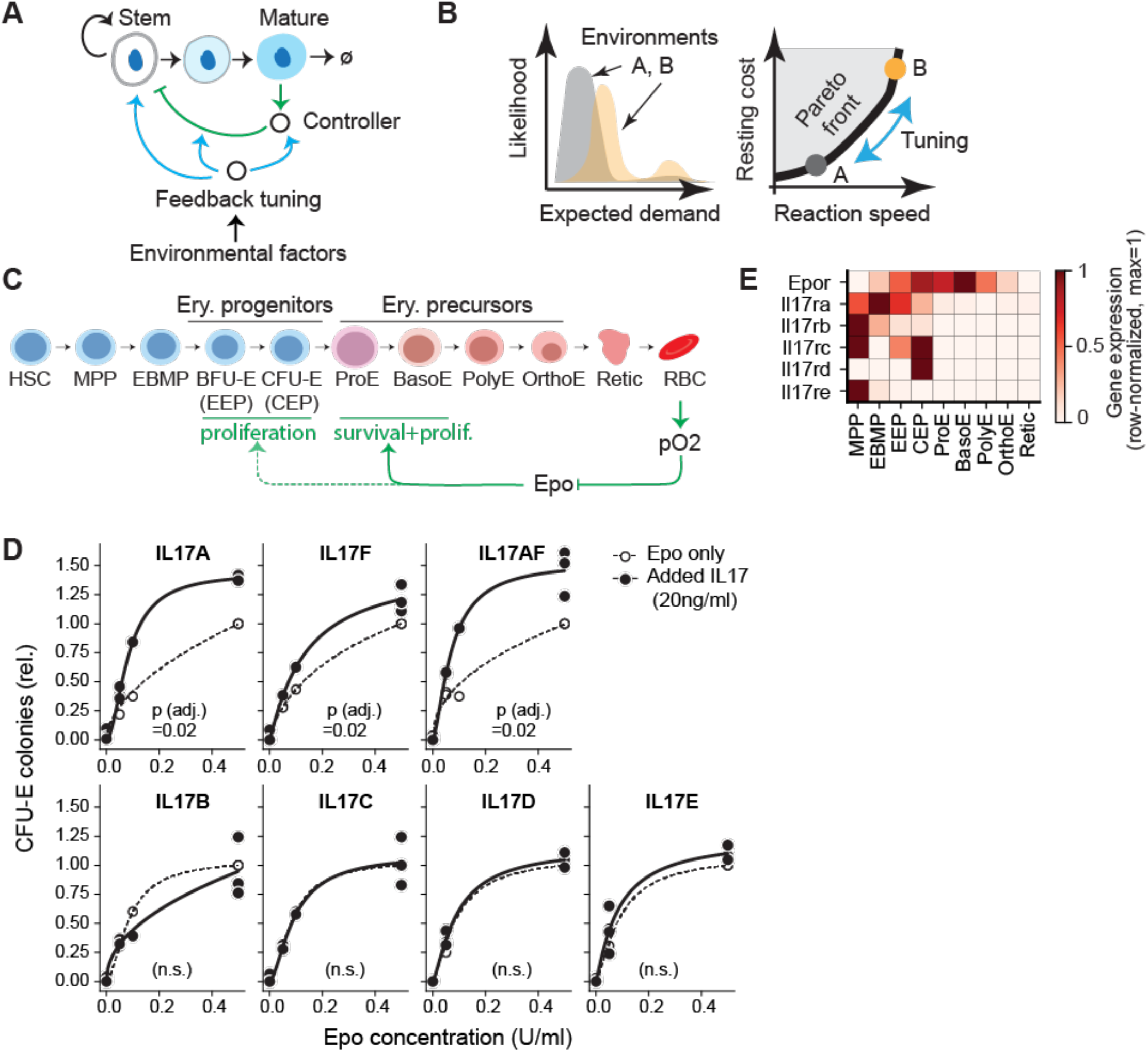
IL-17 ligands with erythropoietic activity in vitro. **A** Illustration of homeostatic control of tissue stem and progenitor cells. Black arrows=cell flux; green arrows=regulatory control; blue arrows=feedback tuning to adapt performance goals. **B** Illustration of performance tuning. Two environments A and B show different likelihoods of increased cell demand (left panel), and accordingly different optimal cost/reaction-speed trade-offs (right panel). Blue arrows: tuning control circuits on the Pareto front alters reaction speed at the lowest possible cost. **C** The erythropoietic negative feedback cell circuit. Abbreviations: HSC, hematopoietic stem cell; MPP, multipotent progenitor; EBMP, erythroid/basophil-mast cell/megakaryocytic progenitor; BFU-E/EEP, erythroid burst-forming unit or early erythroid progenitor; CFU-E/CEP, erythroid colony-forming unit or committed erythroid progenitor; ProE, pro-erythroblast; Baso/Poly/OrthoE, basophilic, poly-, and ortho-chromatic erythroblasts; Retic, reticulocyte; RBC, red blood cell. **D** IL-17A,F and A/F enhance CFU-E colony formation, while IL-17B,C,D and E do not. Mouse bone marrow cells were plated in the presence of Epo at the indicated concentrations, and either in the presence or absence of IL-17 ligands. CFU-E colonies were scored at 72 hours. Data are pooled from n=3 independent experiments for IL-17A, F, AF, D, E and n=2 for IL-17B, C with 3 to 4 technical replicates per experiment. Lines show fits to a Hill function. padj values show Wilcoxon signed-rank test across all Epo concentrations, with Benjamini-Hochberg (BH) correction across all IL-17 ligands. **E** Expression of Epo receptor and candidate tuning IL-17 receptor mRNA transcripts in erythropoietic progenitors. Data derived from single-cell RNA sequencing. Values show row-normalized gene expression, log(1+CP10K)/ log(1+CP10Kmax).

Here we explore an example of tuning in the homeostatic control of a progenitor cell compartment. We focus on erythropoiesis and its regulation by the pro-inflammatory cytokine interleukin 17 (IL-17A). Erythropoiesis is under tight feedback control to maintain optimal tissue oxygen tension. Red cells, which carry oxygen from lungs to tissues, constitute 84% of all body cells ^9^ and turn over every 100 days ^10,11^. Erythropoiesis therefore consumes substantial resources and is vulnerable to stresses such as nutritional deficiencies and inflammation. In addition to optimizing available resources, homeostatic control of erythropoiesis needs to simultaneously maintain red cell number within a narrow range in the steady state, while preserving the capacity for mounting a rapid and quantitatively appropriate stress response when tissue oxygen tension is threatened, as in blood loss or pneumonia ^10-12^. Under stress, erythropoietic rate increases up to 10-fold within a matter of hours. The circuits responsible for this complex regulation are only partly understood.

Erythropoiesis consists of two principal phases ^13,14^ (**Fig. 1C**): an early progenitor phase followed by erythroid terminal differentiation (ETD). In the early progenitor phase, hematopoietic stem cells and multipotent progenitors (MPPs) differentiate into erythroid progenitors, the burst-forming and colony-forming erythroid units (BFU-E and CFU-E), which undergo transit-amplifying divisions. A transcriptional switch then initiates ETD ^13,15^, where erythroid precursors comprising pro-erythroblasts (ProE), early and late erythroblasts, undergo 3-5 maturational cell divisions and enucleate into nascent red cells (reticulocytes). Homeostatic control is established through a lineage-specific and essential feedback mediator, the cytokine hormone erythropoietin (Epo). Epo is secreted by the kidney; its production increases in response to decreasing tissue oxygen tension^16,17^; and it in turn increases erythropoiesis by promoting survival ^18-20^ and cycling^21^ of ProE and early erythroblasts in ETD in a dose-dependent manner. Thus, Epo mediates a closed homeostatic feedback loop (**Fig. 1C**).

Epo also regulates the early progenitor cells prior to ETD, but its regulatory role on these cells is not well explored. MPPs, BFU-E and early CFU-E nonetheless express the Epo receptor (EpoR)^13^ but they do not require Epo for survival^22^ and are less sensitive to Epo stimulation *in vitro* than ETD precursors ^23-25^. In addition to Epo, the early progenitor compartment is also a target for multiple factors with pan-hematopoietic or multi-lineage effects such as stem-cell factor (SCF), BMP4 and glucocorticoids that have established roles in the erythroid stress response^26-29^. Further, our recent single-cell transcriptomic analysis revealed additional growth factor and cytokine receptors expressed in erythroid progenitors ^13^. One such receptor is IL-17RA, whose ligand, IL-17A, is a pro-inflammatory cytokine produced by T cells and other immune cells^30,31^ with pleiotropic tissue targets ^32,33^. We found that *in vitro*, IL-17A enhances the response of both human and mouse CFU-E to Epo^13^. IL-17A is representative of multiple pleiotropic factors in raising three key questions: how does its signaling integrate into a specific and appropriate erythroid output, whether it functionally tunes feedback control, and what control problems might this factor be optimizing for.

Here we investigate the role of IL-17A in erythropoiesis, finding that it has a striking synergistic interaction with Epo *in vivo*, which augments and accelerates the response to hypoxic stress. We show that IL-17A mediates this interaction by increasing the sensitivity of early erythroid progenitors to Epo, an efficient mechanism for tuning Epo-mediated feedback control. Our work suggests potential therapeutic roles for IL-17A in anemia. Further, it offers a template for understanding its action in other cell lineages and suggests a framework for studying how stem and progenitor cells harness broadly-acting environmental cues to tune feedback controls and achieve rapid and coherent lineage-specific responses.

## Results

### IL-17A, IL-17AF, and IL-17F stimulate erythroid colony formation

We recently found that *Il17ra*, encoding an IL-17 receptor chain, is expressed by early hematopoietic and erythroid progenitors^13^. *In vitro* analysis showed that one of its ligands, IL-17A, markedly potentiates formation of CFU-E colonies from adult mouse or human bone marrow in response to stress-levels of Epo (≥0.05 U/ ml), with saturable kinetics and an EC^50^ of ∼60 pM. This effect is lost in cells from *Il17ra*^*-/-*^ mice.

IL-17RA is a common receptor chain in several heteromeric receptor complexes, each of which binds distinct members of the IL-17 cytokine family ^31,32^. We examined each of the seven dimeric IL-17 family ligands for potential erythropoietic activity, using CFU-E colony-formation assays (**Fig. 1D**). We found that the homodimeric IL-17A and IL-17F, and the heterodimeric IL-17AF, all enhanced CFU-E colony formation in response to Epo, though none had an effect in the absence of Epo. By contrast, the remaining homodimeric IL-17 ligands, IL-17B, C, D and E, had no effect. Thus, addition of IL-17A (20ng/ml) to the medium increased the response to Epo (0.5 U/ml) by 39% ± 1.4% (mean ± SEM, adjusted p value p_adj_ = 0.02, Wilcoxon signed-rank test paired by Epo concentration, with Benjamini Hochberg (BH) correction across cytokines). Similarly, addition of IL-17AF increased the maximal response to Epo by 46% ± 1.1% (p_adj._= 0.02) and IL-17F caused an increase of 21% ± 0.7% (p_adj._= 0.02). IL-17A, F and AF are all known to bind the same IL-17RA/IL-17RC receptor complex^31,34^, suggesting that this is the functional IL-17 receptor in erythroid progenitors. Indeed, both these receptor chains are expressed in erythroid progenitors (**Fig. 1E**).

As a technical note, the efficacy of IL-17A, F and AF in CFU-E assays was sensitive to the age of the lyophilized proteins and to the length of storage after reconstitution (**Fig. S1A**). If overlooked, these factors introduce a significant source of variation. The non-responsiveness to IL-17D and E could not however be explained by cytokine age.

### IL-17A and Epo synergize in vivo to increase erythropoietic rate

We next examined whether IL-17A promotes erythropoiesis *in vivo*. We established an optimal dosing regimen by undertaking pharmacokinetic analysis of IL-17A (**Fig. S2A**, Wu et al, in review). We then treated 8 to 12-week-old Balb/c mice with either vehicle (PBS), IL-17A (200ng/g every 12 hours), Epo (0.25U/g every 24 hours), or both Epo and IL-17A (n= 6 to 8 mice per treatment group, pooled from 3 independent experiments). Mice were sacrificed at 72 h, and blood, bone marrow and spleen were collected for a comprehensive analysis of hematopoietic progenitors and mature blood cells (**Fig. 2, Figs. S2-S4**). We first examined spleen size, which, as expected, was increased 2-fold by Epo treatment. IL-17A by itself had little effect, but the combined Epo and IL-17A treatment increased spleen size by nearly 3-fold [**Fig. 2B**; spleen to body weight ratio for vehicle, 0.34±0.008% (mean ± SEM); for Epo, 0.67±0.06%; and for Epo+IL-17, 1.0±0.03%; the difference between Epo and Epo+IL-17A is significant at p=0.0003, 2-tailed *t-*test].

**Figure 2:**
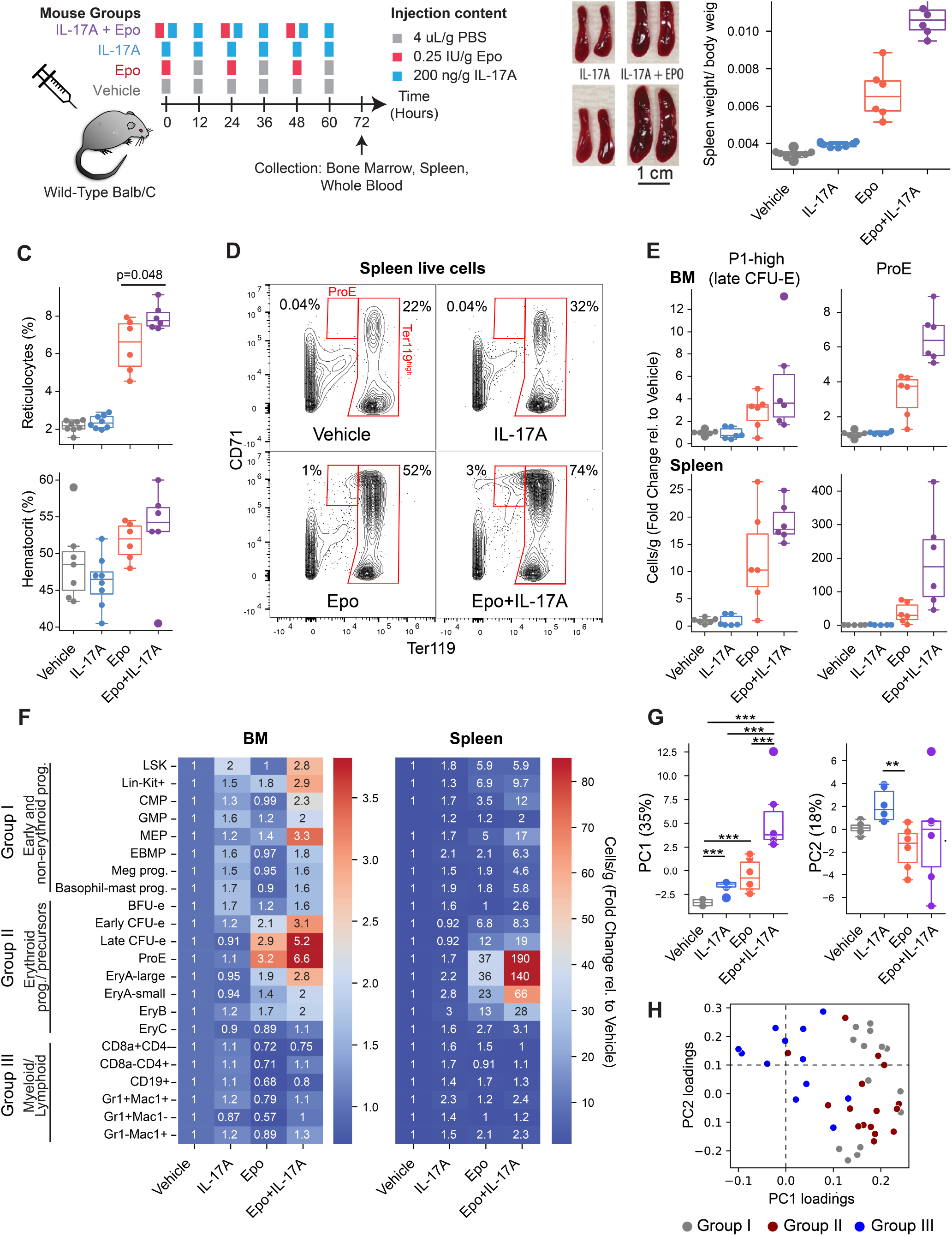
IL-17A synergizes with Epo to increase erythropoietic rate in vivo. **A** Experimental design to evaluate IL-17A erythropoietic stimulation *in vivo*. Resulting data in panels B to G pooled from 3 independent similar experiments, with a total of 6 to 8 mice per treatment. FACS gate definitions and full data are in **Figs. S2** to **S4. B** Representative images of freshly-isolated spleens at 72 hours and spleen to body weight ratios. p-values are from two-tailed t-tests. Here and in all subsequent boxplots, data points correspond to individual mice, box shows medians and the central quartiles (25th to 75th percentiles), and whiskers extend by 1.5 times the inter-quartile range. **C** Changes in blood reticulocyte fraction (CD71^+^ Draq5^-^ blood cells) and hematocrit. **D** Representative flow-cytometric 2D contour plots of Ter119 (erythroid-specific) and CD71 expression, identifying ProE and later erythroid precursors (Ter119^high^ gate). Erythroid precursor are further sub-divided into sequentially more differentiated erythroblast gates EryA, B and C as in^36^, shown in **Fig. S2B. E** Summary data of bone-marrow (BM) and spleen cells labeled with either a 10-color stain antibody-panel to identify CFU-E ^13,37^ (**Fig. S3 A**) or with the CD71/Ter119 panel to identify ProE (panel D). **F** Heat map summarizing the fold-change in the mean cell number (cells/g body weight) of n=6 mice per condition relative to vehicle-treated mice, for each flow-cytometric subset in bone marrow (BM) and spleen. Full data for individual subset/mice is given in panel E, **Figs. S2-S4**, and **Table S1**. Groups I-III indicate functionally distinct progenitor subsets. **G** Principal component (PC) analysis of the unlabeled (mouse x subset) data matrix underlying panel F, treating bone marrow and spleen subsets as independent features, for all mice in the 4 treatment groups. PC1 and PC2 scores for mice grouped by treatment (35% and 18% of the between-mouse variance explained respectively). Statistical significance was assessed with Kruskal-Wallis H-test across all treatments (one-way ANOVA on ranks; p=0.0002 for PC1, p=0.0147 for PC2) and a Conover test was used for post-hoc pairwise comparisons with Benjamini Yekutieli (BY) correction: ***, p_adj_<10^−3^; **, p_adj_<10^−2^. **H** Flow-cytometric subset loadings of PC1 and PC2, colored by the groups defined in panel **F**. PC1 is enriched for early hematopoietic progenitors (Group I) and erythroid progenitors and precursors (Group II; p=1.45 x 10^−6^, Wilcoxon rank sum test comparing PC1 loadings for Groups I and II v. Group III). PC2 is enriched for myeloid and lymphoid subsets (Group III, p=0.029).

Spleen enlargement indicates extra-medullary hematopoiesis. Blood reticulocytes, increased by Epo, were further increased by combined Epo+IL-17A treatment (**Fig. 2C, p=0.048**), suggesting the enlarged spleens reflect expanded erythropoietic tissue and accelerated erythropoiesis. Given that the hematocrit was unaltered in any of the treatment groups (**Fig. 2C**), the observed accelerated erythropoiesis is the direct result of erythropoietic stimulation by the combined Epo+IL-17A treatment, as opposed to a compensatory response to anemia/hypoxia. Of note, 72 hours of erythropoietic stimulation, while increasing reticulocytes, is not expected to result in a significant change in hematocrit.

We next examined early hematopoietic progenitors, erythroid progenitors and precursors, and myeloid and lymphoid cells in the bone marrow and spleen (**Figs. 2D-F, Figs. S2-S4, Table S1**). First, using CD71 and Ter119 antibodies, we found the expected increase in ProE and in Ter119^high^ erythroblast subsets EryA/B/C ^35,36^ in response to Epo in both bone marrow and spleen (**Fig. 2D-F and Fig. S2B**). While there was no response to IL-17A alone, there was a striking, enhanced response to the combined Epo+IL-17A treatment, across all erythroblast subsets, peaking in ProE and EryA: spleen ProE increased 37±12-fold in response to Epo, whereas the combined Epo+IL-17A treatment resulted in a 192±58-fold increase. Second, we examined changes in the abundance of early progenitors by employing a recent flow-cytometric strategy that identifies highly pure BFU-E and CFU-E, megakaryocytic progenitors, basophil-mast cell progenitors, and oligopotent erythroid/megakaryocytic/basophil-mast cell progenitors (EBMPs)^13,37^ (**Fig. S3A**, the ‘10 color stain’). With this approach, we found enhanced response of early and late CFU-E to the combined Epo+IL-17A treatment, as compared with Epo-only treatment, in both bone marrow and spleen (**Fig. 2E-F, Fig. S3C**). And third, we examined changes in abundance of early hematopoietic progenitors: there were enhanced responses of CMP and MEP progenitors to the combined Epo+IL-17A treatment, when compared with Epo alone (**Fig. 2F, Fig. S3E**).

Taken together, these results show that combined Epo+IL-17A treatment elicits an enhanced response from erythroid progenitors and precursors, and from early hematopoietic progenitors that contribute to the erythroid lineage. To formalize this observation, we carried out two statistical tests: first, we divided the examined progenitors into three principal groups and then tested for changes in abundance for each group: early hematopoietic progenitors; erythroid progenitors-and-precursors; and myeloid-and-lymphoid progenitors. These tests confirmed that early hematopoietic progenitors and erythroid progenitors increased following the combined treatment as compared with Epo alone (p_adj._= 0.017 in bone marrow by Wilcoxon signed-rank test paired by FACS gate, with BH correction across comparisons; and p_adj._=0.03 in spleen), while there was no significant effect of the combined treatment on myeloid and lymphoid progenitors (**Fig. 2F, Fig. S2C,D**). Second, we carried out a principal component analysis (PCA) of all (>40) flow-cytometric subsets analyzed in each condition. The first principal component, which explained 35% of the variance between mice from all conditions (**Fig. 2G**), corresponded to an erythroid response as seen from its loading coefficients (**Fig. 2H**). This principal component ordered the four treatment groups with untreated and IL-17A-treated mice corresponding to lower values, followed by Epo-treated and Epo+IL-17A-treated mice (**Fig. 2G**). By contrast, the second principal component (18% variance explained) partitioned IL-17A from the Epo-treated condition and was enriched for myeloid and lymphoid subsets, including GMP progenitors, CD4+ and Gr^+^/Mac1^+^ precursors (**Fig. 2G,H**).

Together, these results show that IL-17A synergizes with Epo: it is an erythropoietic stimulating agent *in vivo*, but only in the context of elevated Epo, as might be seen in erythropoietic stress.

In these experiments we also examined the well-established pro-survival effect of Epo in erythroblasts^19,36^ by staining with Annexin V, which marks apoptotic cells. We found reduced Annexin V binding to erythroblasts in response to both Epo and Epo+IL-17A treatments, but to approximately similar levels (**Fig. S4D**). This suggests that the enhanced response to the combined Epo+IL-17A treatment is mediated through a mechanism distinct from erythroblast survival.

### IL-17A is dispensable for baseline erythropoiesis

The ability of IL-17A to accelerate erythropoiesis only in the presence of elevated Epo suggests it is unlikely to contribute to basal erythropoiesis. To test this, we examined mice with germline deletion of *Il17ra*^38^. Hematocrit, red blood cell and reticulocyte numbers in *Il17ra*^*-/-*^ mice were similar to controls (**Fig. S5A, B**). However, they had significantly enlarged spleens (**Fig S5A**). Flow cytometric analysis showed this to be the result of expansion of spleen ProE and erythroblasts (**Fig S5C**). Together these observations suggest compensated anemia in *Il17ra*^*-/-*^ mice, possibly a result of chronic inflammation. Indeed, the latter is supported by the observed basophilia, possibly a result of impair anti-bacterial and anti-fungal immunity in IL-17A deficiency ^39,40^ (**Fig S5A**).

We therefore proceeded to examine hematopoietic and erythroid-specific deletions of *Il17ra*, crossing a mouse model expressing a ‘floxed’ version of the receptor ^41^ with Vav-iCre ^42^ and EpoR-Cre ^43^ mice. Quantitative-PCR (qPCR) analysis showed poor deletion in EpoR-Cre mice, possibly because *Il17ra* is expressed earlier than *Epor* during erythroid differentiation (**Fig. 1E**). By contrast, *Il17ra* was deleted in over 99% of Kit^+^ and in ProE cells on the Vav-iCre background, in both bone marrow and spleen (**Fig. 3A, Fig S5D**). Unlike the germline deletion, there was no increase in spleen size in the IL-17ra^f/f^/Vav-iCre mice (**Fig. 3B**) and no expansion of splenic erythroid populations (**Fig. S5E**), confirming that the enlarged spleen we observed in the *Il17ra* germline deletion is the result of non-cell autonomous effects such as chronic infection. The hematocrit in the IL-17ra^f/f^/Vav-iCre mice was normal **(Fig 3B**). We conclude that there is no requirement for IL-17RA in the maintenance of basal erythropoiesis.

**Figure 3:**
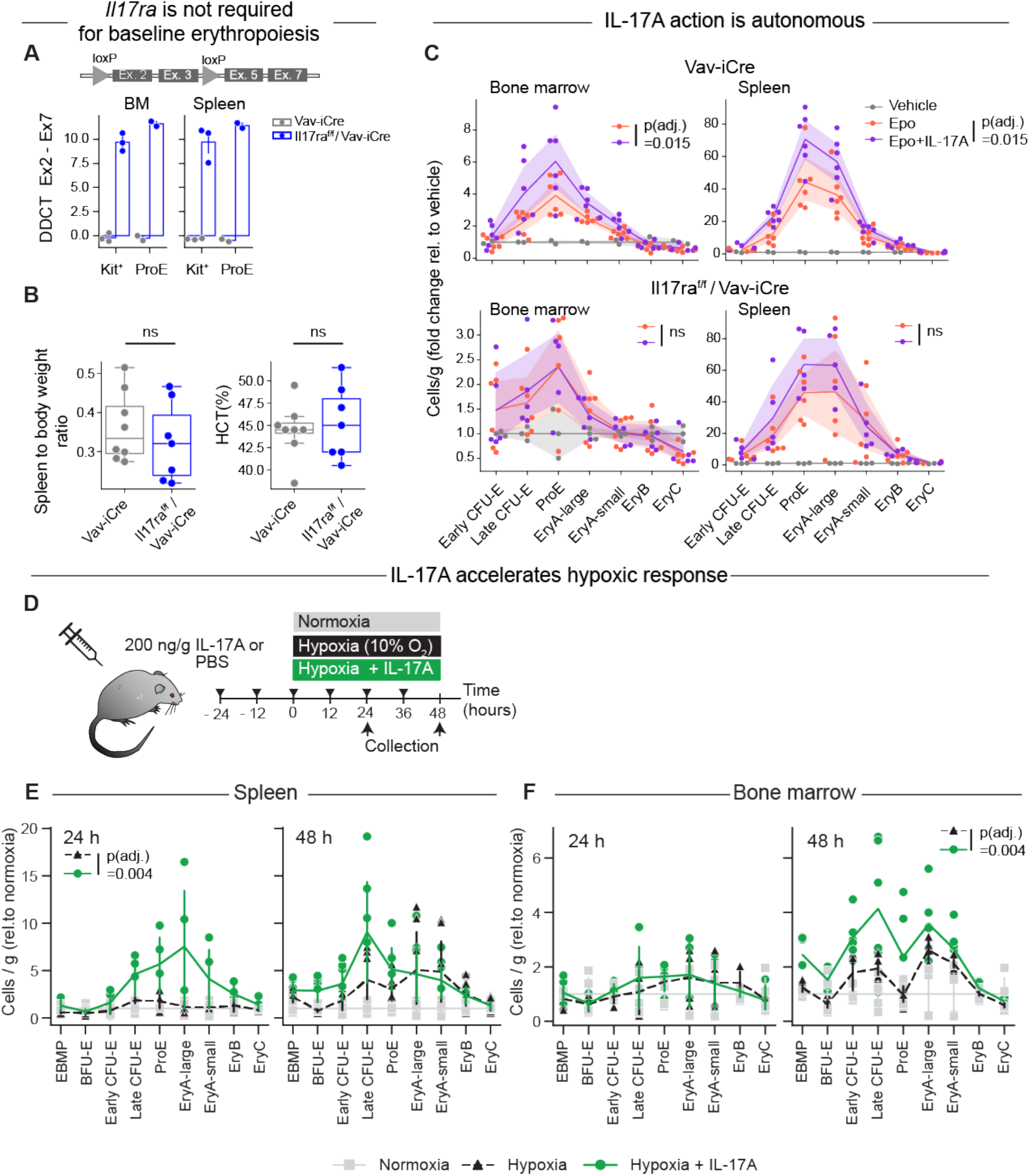
The erythropoietic effect of IL-17A is autonomous to hematopoietic cells and accelerates the response to hypoxic stress. **A** Deletion efficiency of *Il17ra* in the hematopoietic-specific Vav-iCre model, evaluated by genomic qPCR of the excised genomic locus (exons 2,3) normalized to an adjacent locus (exons 5,7). The cartoon of the *Il17ra* locus is not to scale. qPCR was performed on FACS-sorted Kit^+^ and ProE cells. Data points are independent biological replicates. Additional analysis confirming deletion using independent primers is in **Fig. S5D. B** The IL-17RA receptor is not required for steady-state erythropoiesis as seen from the stable baseline (uninjected) spleen to body weight ratio and hematocrit in Il17ra^f/f^ / Vav-iCre mice. Consistent results are seen in analysis of hematopoietic subsets by flow cytometry in **Fig. S5E. C** Epo/IL-17A synergy requires IL-17RA expression in hematopoietic cells, as seen by comparing erythroid progenitor cell numbers at 72 hours following cytokine injections in control Vav-iCre and Il17ra^f/f^ / Vav-iCre mice. Cytokine injections followed the schedule in **Fig. 2A**. Data are pooled from 2 independent experiments, n=6 mice per treatment. Color bands are 95% confidence intervals. p_adj_ values are one-tailed Wilcoxon signed-rank test paired by progenitor subset and then BH-corrected across the hypotheses, testing whether Epo+IL-17A results in larger progenitor populations than Epo alone. **D** Experimental design for evaluating IL-17A tuning a hypoxic response. Triangles indicate injections of IL-17A or PBS. **E,F** Flow-cytometric subsets in spleen and bone marrow respectively at the indicated time points during hypoxia. Results are expressed as a fold-change in the viable cells per gram body weight, relative to equivalent progenitor subsets in normoxia. Data pooled from four independent experiments (2 experiments each at 24 h and 48 h), for a total of n=6 mice for each time point for hypoxia±IL-17A and n=4 mice for normoxia. Datapoints are individual mice, lines are drawn through the means. P-values are Wilcoxon signed-rank test across all erythroid subsets at each time point and tissue, testing that erythroid subsets are larger in hypoxia +IL-17A than in hypoxia. BH correction was applied across comparisons.

### Synergism between IL-17A and Epo is autonomous to hematopoietic cells

Since the synergism between Epo and IL-17A is observed in CFU-E assays *in vitro* (**Fig. 1D**), it is likely to be cell autonomous. To investigate this *in vivo*, we tested whether synergism persists in the context of a hematopoietic-specific *Il17ra* deletion. We injected IL-17ra^f/f^/Vav-iCre and control Vav-iCre mice with either Vehicle, IL-17A, Epo or Epo+IL-17A for 72 hours, following the same experimental design we previously used in wild-type mice (n=6 mice per treatment/genotype combination). We found no significant difference between Epo and the combined Epo+IL-17A treatment in IL-17ra^f/f^/Vav-iCre mice, while synergism was still observed in the response of both spleen and bone-marrow erythroid progenitors and precursors in control Vav-iCre mice (p_adj_.= 0.015, **Fig. 3C**). We therefore conclude that the synergistic interaction between Epo and IL-17A is autonomous to the hematopoietic system.

### IL-17A accelerates the erythropoietic response to hypoxia

Tissue hypoxia induces an erythropoietic stress response, driven by increased Epo secretion from the kidney. The resulting higher circulating Epo concentration accelerates erythropoietic rate by promoting survival and cycling of erythroid precursors (**Fig. 1C**). We asked whether IL-17A could stimulate erythropoiesis in the context of physiological stress, without administration of exogenous Epo.

We previously modeled erythropoietic stress by placing mice in a reduced-oxygen environment (10% O_2_, compared with 21% oxygen at sea level), finding that blood Epo levels increase and plateau at 0.03 U/ml within 24 hours of the onset of hypoxia, increasing ProE and EryA in spleen and bone marrow ^19^. To test whether IL-17A impacts the hypoxic stress response, we injected mice with either IL-17A (200 ng/g every 12 hours) or vehicle for 24 hours before either transferring them to a hypoxic environment for up to 48 hours or allowing them to remain in normoxia. We continued IL-17A administration up until the end of the experiment (**Fig. 3D**).

Older reports based on colony formation assays show that in response to acute erythroid stress, bone-marrow erythroid expansion is delayed relative to the spleen, since bone-marrow BFU-E initially migrate to the spleen, where they contribute to rapid erythroid expansion ^44-46^. Here, our flow-cytometric assessment of BFU-E and CFU-E using the ‘10 color stain’ ^13,37^ (**Fig. S3A**) is in agreement with these older studies, showing an initial decline in bone-marrow BFU-E and a slower bone-marrow erythroid expansion relative to the spleen (**Fig 3E, F and Table S2**).

Significantly, treatment with IL-17A accelerated the erythroid response in both tissues. At 24 hours in hypoxia, IL-17A treatment increased nearly all of the spleen progenitors and precursors on the erythroid differentiation trajectory when compared with hypoxia alone (early and late CFU-E, ProE, and EryA/B/C, p_adj._= 0.004, Wilcoxon signed-rank paired by cell subset, with BH correction) (**Fig. 3E**). As an example, splenic ProE increased by 1.76±0.48 fold in hypoxia, but by 5.7±1.9 fold in combined hypoxia with IL-17A. In the bone marrow, the response to hypoxia became apparent at 48 hours, with IL-17A treatment further increasing erythroid progenitors and precursors (**Fig. 3F**, p_adj._= 0.004, test as above). Late CFU-E, for example, increased by 1.9±0.15 fold in response to hypoxia, but by 4.1±1.0 in response to hypoxia with IL-17A. Taken together, these findings show that IL-17A accelerates the erythropoietic response to hypoxia.

### Hypotheses for the function of IL-17A in feedback control: 1. IL-17A as a hypoxia signal

The work presented above shows that IL-17A enhances the erythropoietic stress response *in vivo*, and that it does so through a synergistic interaction with Epo. This finding led us to two related questions: first, what physiological function does IL-17A fulfill in erythropoietic feedback control? And second, by what mechanism does IL-17A exert the observed synergy with Epo?

Tissue injury and inflammation may lead to hypoxic tissue microenvironments^47^, and vice versa, tissue hypoxia may lead to inflammation^48^. IL-17A has been implicated in both these settings^49^. We therefore considered two possible complementary hypotheses for the role of IL-17A: first, IL-17A might act as a hypoxia signal, providing a second feedback pathway independent of Epo and the kidney. Second, IL-17A produced independently of hypoxia, for example in response to infections, may signal increased risk of hypoxia in the near future. Under this alternative hypothesis, IL-17A pre-emptively tunes the response to Epo to achieve faster recovery should hypoxia occur.

We first considered the possibility that IL-17A may serve as a second hypoxia signal. Such a signal may help erythroid progenitors to disambiguate the onset of low levels of hypoxic stress, by providing two independent signals that indicate hypoxia. Motivation for this hypothesis comes from prior work that shows induction of IL-17A by HIF1 in T cells ^48,50^, and elevated IL-17A in response to high-altitude hypoxia ^51^.

To explore whether this hypothesis is plausible, we asked whether IL-17A increases in response to hypoxia. We found that mice placed in hypoxia (10% oxygen) for 24 hours, had a ∼five-fold increase in plasma IL-17A compared to normoxic mice (5.49 ± 2.09 pg/mL vs 1.19 ± 0.52 pg/mL, p=0.03, n= 31 mice) (**Fig. S6A**). This is similar to the ∼3 fold increase in plasma Epo in response to the same degree and length of hypoxia ^19^ but is at least 300-fold lower than the EC_50_ for the erythroid stimulatory effect of IL-17A (2ng/mL) in CFU-E colony formation *in vitro* ^13^. Several estimates of baseline blood plasma IL-17A concentrations in humans are higher than our measured value here in mice but are still at least 10-fold lower than the EC_50_ for IL-17A in CFU-E assays (summarized in Table S3). Thus, IL-17A is unlikely to function as a second hypoxia signal in the feedback control of erythropoiesis, though we cannot fully rule out this hypothesis without confidence in endogenous IL-17A activity following hypoxia. We did not pursue this hypothesis further.

### Hypotheses for the function of IL-17A in feedback control: 2. A model of erythropoietic speed/burden tuning by IL-17A

We next considered a model where IL-17A serves to tune erythropoiesis in anticipation of future hypoxic stress. Evidence for the feasibility of this hypothesis comes from work that finds increasing levels of IL-17A preceding hypoxic stress in infectious pneumonia. As examples, IL-17A increases early in experimental pneumococcal pneumonia, and is an early marker of likely disease severity and development of acute respiratory distress syndrome in Covid-19 patients ^52-54^. Levels of IL-17A in patients hospitalized with viral infections, including SARS-CoV2 or MERS, increase 3 to 20 fold over baseline levels ^54-57^ and can exceed the EC_50_ for the erythroid stimulatory effect of IL-17A *in vitro* (2ng/mL) (Table S3).

Tuning feedback by IL-17A is premised on the idea that the erythropoietic system faces a trade-off between its capacity for rapid response, and a burden imposed by over-production of erythroid progenitors. ProEs are produced in excess in the steady-state and undergo apoptosis in the absence of a strong pro-survival Epo signal ^18-20,36,58^. Our finding here that IL-17A accelerates the response to hypoxic stress (**Fig. 3E,F**) suggests that it might do so by tuning some parameters of the erythropoietic feedback control circuit, potentially changing the constitutive response-burden trade-off. The trade-off may favor a lower burden in health, but a faster response and a higher burden when hypoxic risk is elevated (**Fig. 1B**).

To ground this idea and guide interpretation of experimental results, we developed a simple mathematical model (**Fig. S6B,C** and Supplemental Text 1), shown schematically in **Fig. 4A**. Here, (i) MPPs give rise to transit-amplifying erythroid progenitors (BFU-E and CFU-Es); these give rise to precursors (ProE), that differentiate, ultimately giving rise to mature RBCs; (ii) RBCs carry oxygen from lung to tissue; (iii) tissue pO_2_ represses Epo production; and (iv) Epo levels determine ProE survival, a well-established mechanism through which Epo regulates erythropoietic rate ^18-20,36,58^. Finally, (v) expansion of early erythroid progenitors may be regulated through both Epo-dependent and Epo-independent pathways. In agreement with older work, BFU-E and CFU-E are independent of Epo for survival and are relatively insensitive to Epo-mediated expansion in the absence of stress (healthy individuals in normoxia) ^10,19,59^.

**Figure 4.**
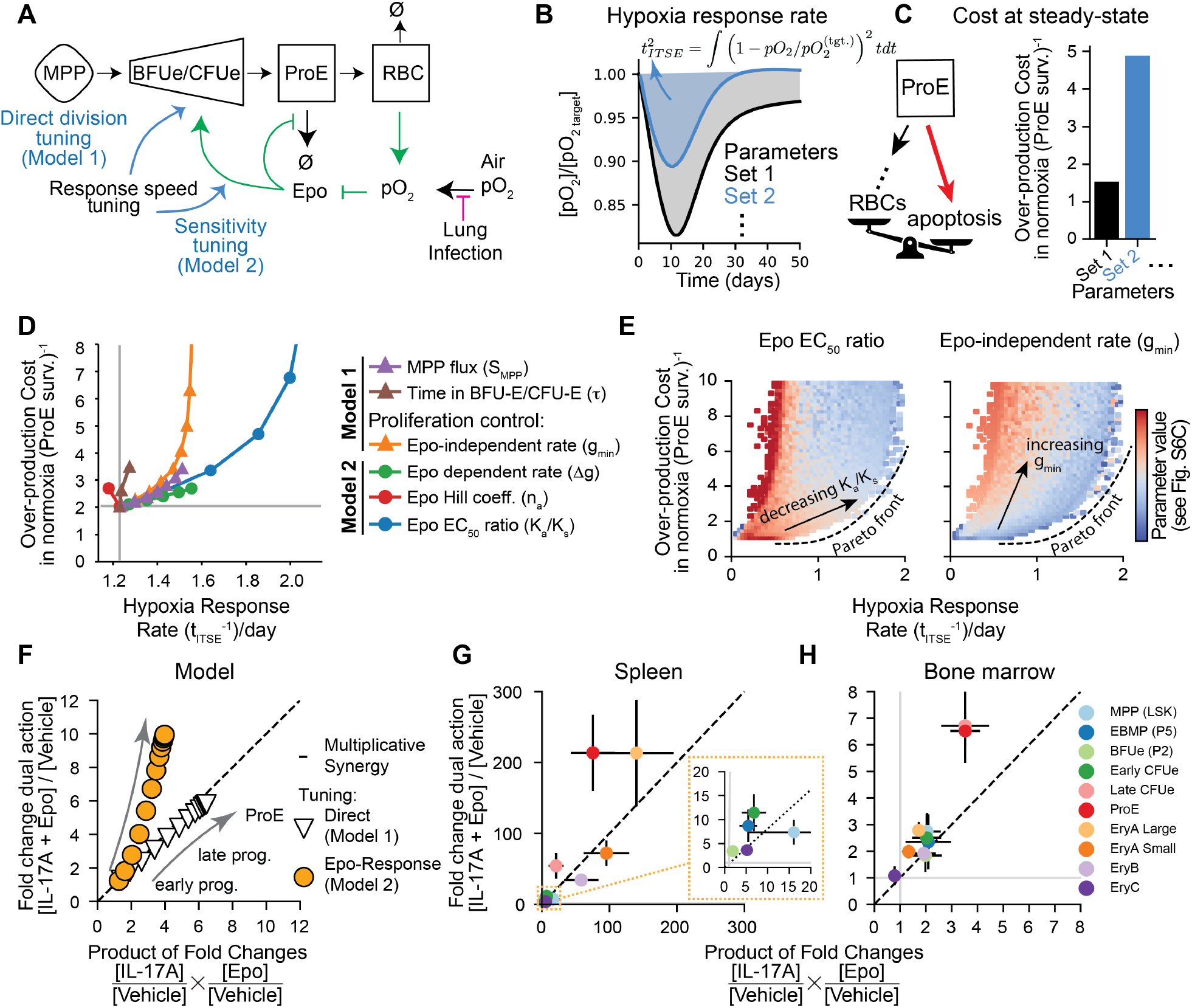
A dynamical model predicts optimal tuning mechanisms for erythroid feedback circuit. **A** Schematic of a simplified dynamical systems model for evaluating response-speed/burden trade-offs in erythropoiesis. Full model parameterization is in **Fig. S6B** and **Supplementary Text 1**. Mechanisms for tuning by IL-17A (blue arrows) can be summarized into two classes: Model 1 (tuning progenitor flux directly) and Model 2 (tuning Epo sensitivity of progenitors). **B** Illustrative numerical solutions of the dynamical model for pO_2_ blood concentration following gradual hypoxic onset. Two different configurations of the system (Parameter Sets 1,2) show different recovery times (defined as the root integral time squared error (*t*_*ITSE*_) of the pO_2_ levels from target). **C** The constitutive over-production burden during normoxia for the same parameter sets as in (**B**). The over-production burden results from high levels of apoptosis of ProE (shown schematically), and is defined as 1/(ProE survival). The faster response in (**B**) requires a higher burden. **D** Tuning individual control parameters from a baseline state identifies variation in their contribution to recovery speed (*t*_ITSE_)^-1^ after pO_2_ perturbation and the over-production burden [defined in (**C**)]. Parameters corresponding to models 1 and 2 (panel A) are denoted by triangles and circles respectively. **E** Random sampling control parameters over physiological ranges reveals a Pareto front in the response/burden trade-off. The heatmaps shows the values of two parameters representative of Model 1 (Epo-independent proliferation, *g*_*min*_) and Model 2 (Epo-dependent proliferation EC_50_, *K*_*a*_). Color bar units and additional parameter values are given in **Fig. S6F. F** Computational predictions for the expansion of erythroid progenitors in each of the two models following Epo + IL-17A treatment as compared to multiplicative fold change of Epo and IL-17A alone. The dashed line indicates **Figure 4 (Continued)** multiplicative synergy. Model 1 is implemented by modulating *g*_*min*_ while model 2 is implemented by modulating *K*_*a*_ (this figure) or Δ*g* (**Fig. S6G**). Refer to **Supplementary Text 1** for modeling parameters used. **G,H** Measured fold changes in cell populations from spleen (**G**) and bone marrow (**H**) comparing [Epo + IL-17A]/[Vehicle] treatment to the expected multiplicative fold change ([IL-17A]/[Vehicle] × [Epo]/[Vehicle]). Data points above the dashed line indicate above-multiplicative synergy. Inset in (**G**) shows zoomed view of lower fold changes. The same experimental data as in **Fig. 2F**.

We investigated cost-performance trade-offs in this model using numerical simulations to test how different strategies for tuning the erythroid feedback circuit might alter the speed of recovery following a drop in oxygen delivery to tissues, as may occur during respiratory distress. Recovery requires increased RBC production, which increases the number of circulating RBCs, thereby compensating for the reduced oxygen loading of each individual RBC (**Fig. 4B**, hypoxic stress dynamics in **Fig. S6D**). Upon tuning the model, the response rate changes (quantified by the integral-time-squared error or ITSE, **Fig. 4B**), as does the burden of progenitor over-production during normoxia (**Fig. 4C**). Significantly, the model shows that the continuously dying and regenerating progenitor and precursor reserve is essential for a fast erythropoietic response to sudden hypoxic stress (**Fig. S6E**). It thus recapitulates the experimental observation that a loss of excess production capacity, by genetic elimination of the apoptotic mechanisms that preserve it, delays production of new RBC in response to hypoxia ^20^.

What then are the strategies by which IL-17A might tune homeostatic control? A scan of model parameters in which they are varied one at a time (**Fig. 4D**) reveals only two principal tuning mechanisms modulating the speed/burden trade-off: IL-17A may either directly regulate early progenitor proliferation or their generation by multipotent cells (**Fig. 4A, Model 1** and **Fig. S6B,C**), or it may alter the response of early progenitors to Epo proliferative signals, by altering either Epo’s EC_50_ or the maximal Epo proliferative response (**Fig. 4A, Model 2** and **Fig. S6B,C**). Thus, the model suggests that early progenitors may serve as key targets for response/burden tuning of the erythropoietic system (**Fig. 4A**). These cells are also the likely cellular targets of IL-17A *in vivo*, based on their expression of *Il17ra* ^13^ (**Fig. 1E**), their response to treatment (**Fig. 2**), and our finding that the combined IL-17A and Epo treatment has little further effect on ProE survival beyond that of Epo alone (**Fig. S4E**), suggesting ProE are not direct targets of IL-17A.

The response/burden trade-off for single parameters (**Fig. 4D**) suggested, however, that the two models are not equivalent in their trade-offs. Varying the maximal proliferative response to Epo or varying its EC_50_ (Model 2) provided gains in performance with a smaller associated increase in ProE over-production when compared with varying Epo-independent progenitor proliferation or progenitor production from MPPs (Model 1) (**Fig. 4D**). To evaluate whether these differences held for all model configurations, we carried out additional (10^5^) simulations, each randomly sampling model parameter values, and evaluated the two performance goals in each case. This analysis revealed a clear pareto front in cost-performance trade-off (**Fig. 4E, S6F**). Significantly, parameters associated with Model 2 (Epo sensitization) tuned the feedback circuit along a pareto front where response rates accelerated with the lowest additional burden, while the parameter associated with Model 1 (control of Epo-independent progenitor proliferation) did not. Thus, within our model, tuning the sensitivity of early progenitors to Epo offers the lowest possible progenitor over-production burden for a given speed of hypoxic response.

### Predictions that distinguish direct action by IL-17A (Model 1) from sensitization to Epo (Model 2)

We asked whether experimental evidence distinguishes between IL-17A acting via one model (Model 1) or another (Model 2) (**Fig. 4A,D**). Model 1 assumes that IL-17A directly stimulates expansion of early progenitors, even in the absence of high Epo. By contrast, IL-17A-mediated sensitization of progenitors to Epo (Model 2) predicts only a mild response to IL-17A in the absence of high Epo or stress (**Fig. 4A**). Significantly, the response *in vivo* to IL-17A is consistent with Epo-sensitization (Model 2) and not with direct stimulation of progenitors (Model 1), since there is little response of early progenitors to IL-17A alone (**Fig. 2**).

Agreement of our experimental results with Model 2 and not Model 1 can be further appreciated by quantitatively comparing the observed increase in progenitor pool size in response to the combined Epo + IL-17A treatment, with the expected increase, computed from the response to treatment with each cytokine in isolation. Simulations of Model 1 (Epo-independent action) predict multiplicative synergy for the combined Epo and IL-17A treatment, with the increase in progenitors being the product of the response to each cytokine alone (**Fig. 4F**, on-diagonal). By contrast, Model 2 predicts that expansion of erythroid progenitors and precursors should be consistently larger than expected from the product of their independent action (**Fig. 4F** and **Fig. S6G**, above the diagonal). Analysis of our experimental results show they are in agreement with Model 2 (sensitization to Epo by IL-17A, **Fig. 4G, H**).

To further explore the mechanisms underpinning IL-17A action, we investigated two additional properties differentiating the models. First, the expected transcriptional response of erythroid progenitors should differ between Models 1 and 2. If IL-17A sensitizes early progenitors to Epo (Model 2), this may be evident as amplified induction of EpoR and Stat5 gene targets, similar to findings in the interactions of IL-17A with other cytokines^60-63^. By contrast, Epo-independent action of IL-17A on early progenitors (Model 1) is expected to result in genes that are induced independently of Epo and that might be unique to IL-17A signaling. Second, Epo-sensitization (Model 2) assumes that IL-17A stimulates progenitor cell cycling only in the context of elevated Epo. By contrast, Model 1 assumes that IL-17A directly stimulates erythroid progenitor cycling. In the next result sections, we test these predictions.

### IL-17A amplifies the transcriptional response to Epo

To determine the transcriptional response underlying the synergistic action of Epo and IL-17A, we performed single cell RNA sequencing (scRNA-seq) of bone-marrow and splenic hematopoietic cells from mice treated with Vehicle, Epo alone, IL-17A alone, or combined Epo and IL-17A (n=2 per injection treatment) (**Fig. 5A**) using the same dosing and frequency regimen as in **Fig. 2A**. We ensured representation of rarer earlier progenitor populations alongside more abundant maturing precursors by enriching for Kit^+^ progenitors, and then pooling these with 10% unenriched cells. We analyzed 28,681 cells after filtering and sample demultiplexing, from both spleen and bone marrow (n=2 mice per condition; cell counts in Table S4).

**Figure 5.**
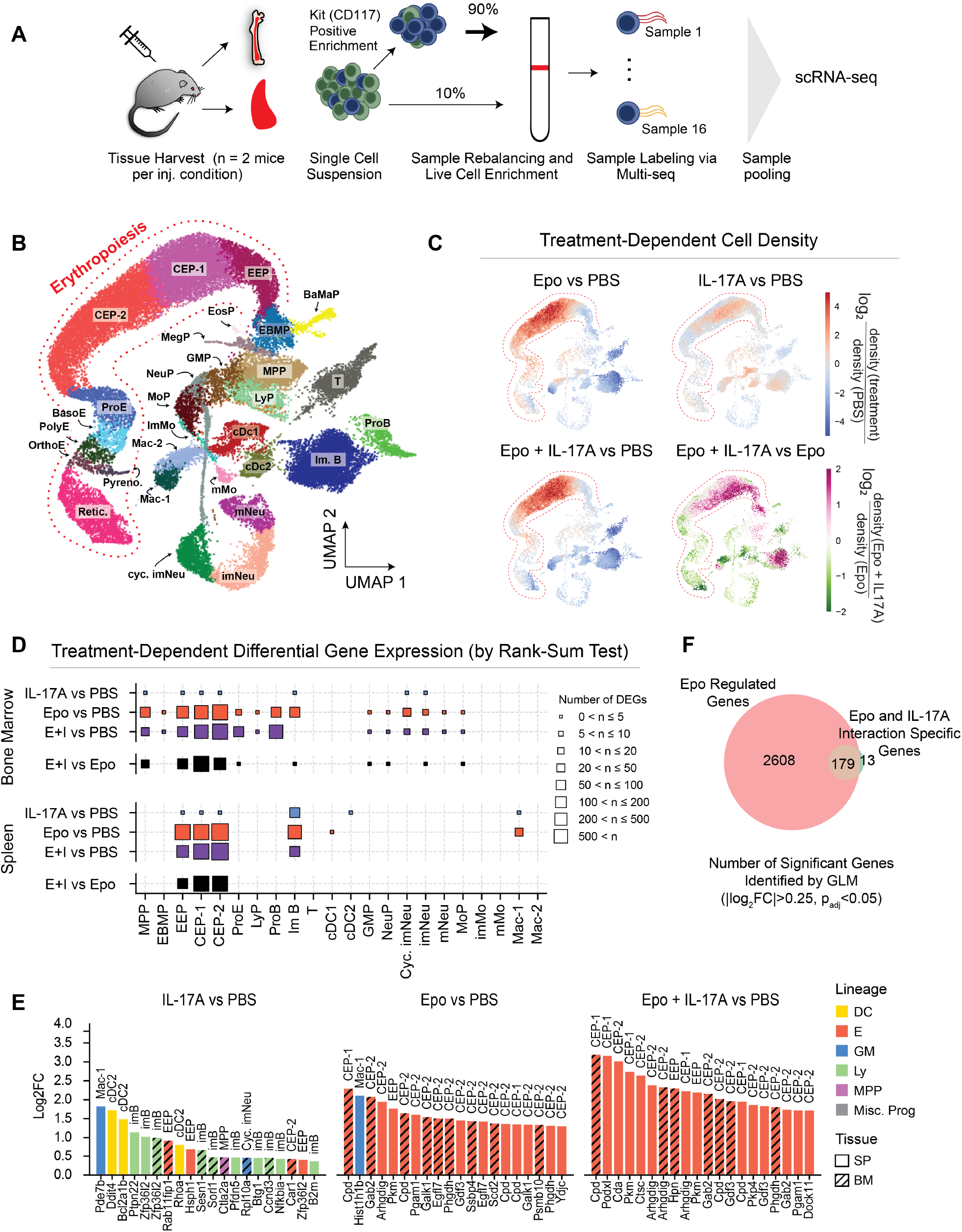
IL-17A scRNAseq analysis reveals a weak response to IL-17A treatment in the absence of Epo. **A** Experimental workflow for scRNA-seq. Bone marrow and spleen samples were harvested (n = 2 mice per treatment), enriched for Kit (CD117)-positive cells, labeled using MULTI-seq, and pooled for sequencing. **B** UMAP visualization of single-cell transcriptomes across conditions and tissues, colored by annotated cell states. MPP, multipotent progenitor; EBMP, erythroid-basophilic-megakaryocytic progenitor; BaMaP, basophil-mast cell progenitor; MegP, megakaryocyte progenitor; EosP, eosinophil progenitor; LyP, lymphoid progenitor; GMP, granulocyte monocyte progenitor; MoP, monocyte progenitor; imMo, immature monocyte; mMo, mature monocyte;NeuP, neutrophil progenitor; imNeu, immature neutrophil; mNeu, mature neutrophil; imB, immature B cell; proB, proB cell; T, T cell; cDC1/2, conventional dendritic cell 1/2; pDC, plasmacytoid dendritic cell; Mac, macrophage; EEP, early erythroid progenitor (BFU-e equivalent); CEP, committed erythroid progenitor (CFU-e equivalent); ProE, proerythroblast; BasoE, basophilic erythroblast; PolyE, polychromatic erythroblast; OrthoE, orthochromatic erythroblast; pyrenocytes, and reticulocytes. **C** Relative cell densities across the sampled RNA-Seq state space, evaluated in high dimensions and visualized on the UMAP plot. The plots are colored by the log2-transformed ratio of cell densities in the neighborhood of each sampled transcriptome, for each treatment pair as indicated. **D** Number of differentially-expressed genes (DEGs) between cells classified to each indicated cell state across treatment conditions in bone marrow and spleen, showing few DEGs in response to IL-17A treatment in isolation, but substantial response to Epo + IL-17A (E+I) as compared to Epo. DEGs are identified by a rank-sum test with BH correction, and |log2(fold-change)|>0.25 and p_adj_<0.05. **E** The top 20 DEGs across conditions and cell populations by magnitude of fold-change reveal weak responses to IL-17A with low lineage-specificity and a robust response in the presence of Epo. Bar height shows log2 fold change; colors indicate lineages: dendritic (DC), erythroid (E), granulocyte-monocyte (GM), lymphoid (Ly), multipotent progenitors (MPP), miscellaneous progenitors (Misc. Prog.). Solid bars represent spleen (SP), hashed bars represent bone marrow (BM). **F** Venn diagram showing the shared identity of genes found to be significantly regulated in a generalized linear model (GLM) that accounts for separate effects of tissue-of-origin, cytokine treatment (Epo or IL-17A), and cytokine interactions (Epo and IL-17A). The overlap of Epo regulated genes and Epo-IL-17A interaction-specific genes show a large proportion of the Epo and IL-17A interaction genes are regulated by Epo alone. See **Fig. S7B,C** for GLM definition and underlying results.

UMAP embedding revealed MPPs, from which emerge nine hematopoietic lineages through a differentiation continuum, consistent with previous studies^13,64-66^ (**Fig. 5B**: erythroid (Ery), megakaryocytic (Meg), basophilic and mast (BaMa), eosinophilic (Eo), lymphoid (Ly), monocytic (Mo), neutrophilic (Neu), and dendritic (DC). For the erythroid lineage, we identified progressive stages of differentiation by transfer learning from reference datasets^13^ where stages were FACS sorted and functionally validated prior to scRNA-Seq. For the remaining lineages we carried out manual curation with lineage-specific marker genes (**Table S4, Fig. S7A**). This data set contains a representation of the complete erythroid trajectory, from MPP to reticulocytes. We note that it strikingly captures differentiation through to erythroblast enucleation, forming pyrenocytes (plasma-membrane-bound nuclei) and enucleated reticulocytes (**Fig. 5B**). This data set allows us to investigate the effects of Epo and IL-17A stimulation across the entirety of the erythroid lineage.

We confirmed that the scRNA-Seq data recapitulates the expected changes in cell type frequency in response to cytokine stimulation by calculating the density of cells along the erythroid trajectory. Density estimation was done in high dimensions and then visualized on the UMAP (**Fig. 5C**). The changes in density were consistent with an Epo- and Epo+IL-17A-dependent expansion of cells classified as early erythroid progenitors (EEP) and committed erythroid progenitors (CEP), which correspond to BFU-Es and CFU-Es respectively^13^. The data cannot reveal absolute changes in cell number due to the cell enrichment strategy used; nevertheless, the density changes are consistent with those seen by FACS (**Fig. 2**), building confidence in analysis of signaling gene expression responses.

The data reveal transcriptional responses of both erythroid and non-erythroid cells to IL-17A. We analyzed both the number of differentially expressed genes (DEGs) and the magnitude of differential expression per gene, comparing each treatment with vehicle, and also comparing Epo alone vs. combined Epo + IL-17A (**Fig. 5D**, showing significant DEGs with |log_2_FC|>0.25). Epo and the combined Epo+IL-17A treatments showed changes across hundreds of genes, with the largest number of DEGs in CEPs (functionally CFU-E), a population that shows overlap in expression of *Epor* and *Il17ra* (**Fig. 1E**). The combined Epo+IL-17A treatment induced multiple DEGs when compared to Epo alone, but by contrast, IL-17A alone induced few DEGs when compared to Vehicle. These results support the view that IL-17A stimulates few changes when acting alone, but is capable of eliciting a response jointly with Epo.

We next examined the identity of the genes responding to Epo+IL-17A treatment. Significantly, the genes responsive to Epo alone were also differentially expressed in response to the combined Epo+IL-17A treatment (**Fig. 5E**), but the magnitude of response was consistently higher in the combined treatment, by up to two-fold. Of the top DEGs (**Fig. 5E**), most have been previously reported as Epo-responsive genes, including *Podxl, Cda, Cpd, Gdfi3, Gab2*, and *Arhgdig*^13,67,68^. To formally show the overlap in the responding genes, we fit gene expression to a generalized linear model that regresses the contribution of the two cytokines, as well as their interactions, and the tissue-of-origin of each cell (spleen or bone marrow; model defined in **Fig. S7B**). This model identified 192 genes whose expression was significantly altered by IL-17A/Epo interaction in erythroid progenitors (|log_2_FC|>0.25, p_adj_<0.05), and significantly 179 of these genes were also regulated by Epo alone (**Fig. 5F, Fig. S7C**).

To quantify the extent to which the combined Epo+IL-17A treatment amplified the response to Epo, we defined an aggregate expression score for an Epo-responsive gene set identified previously^13^ (Table S4). Expression of this gene set was higher in Epo+IL-17A compared to Epo in erythroid progenitors, both in spleen and bone marrow (**Fig. 6A,B**). These changes were not due to global changes in mRNA composition, as can be appreciated by comparison to an expression-matched random gene set, which showed no change in expression in response to the combined treatment (**Figs. S7D**).

**Figure 6.**
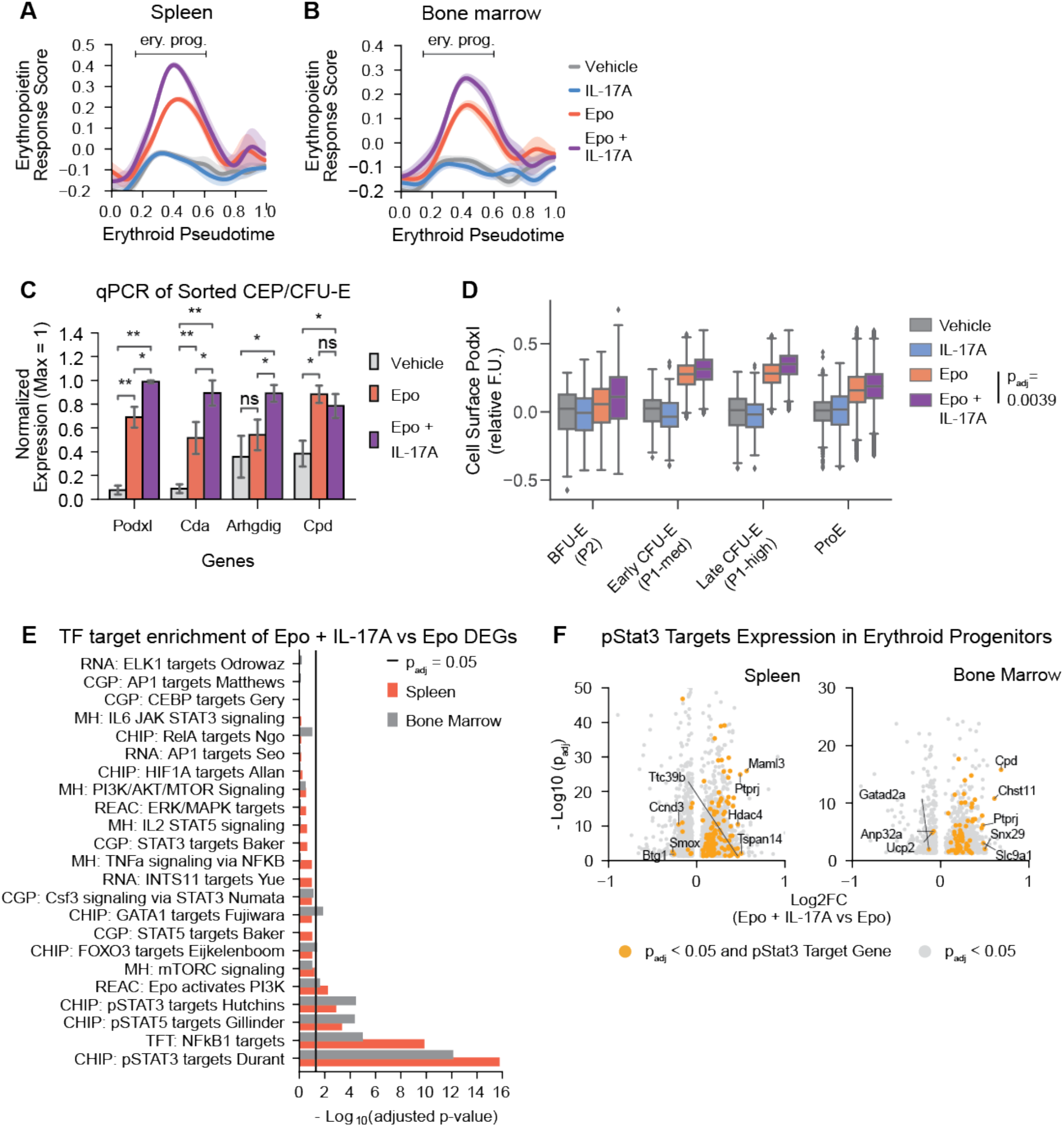
IL-17A amplifies the gene expression response to Epo in erythroid progenitors. **A,B** Erythropoietin response scores evaluated from the scRNA-Seq data (**Fig. 5**) across erythroid pseudotime for different treatment conditions in (A) Spleen and (B) Bone Marrow. The span of time representing erythroid progenitors (EEP and CEP) is marked along the pseudotime. The score is calculated from expression of Epo-responsive genes. **C** Independent evaluation of synergy in expression of selected Epo-responsive gene transcripts (*Podxl, Cda, Arhgdig, Cpd*) measured by qPCR in sorted CFU-E/CEP cells under different treatments (n = 4 mice per treatment, pooled from 4 independent experiments). Error bars represent the standard error. Statistical significance was tested with a one-tailed Wilcoxon rank-sum test *p < 0.05, **p < 0.01, ns: not significant. **D** Podxl cell-surface protein expression measured by flow cytometric in early erythroid progenitors for each of the four treatments. Data pooled from 4 independent experiments. For early progenitors, data is mean of n=3 mice per treatment; for ProE, n=5 mice per treatment. Statistical significance tested with a one-tailed Wilcoxon signed-rank test across FACS subsets. **Figure 2.6 (Continued) E** Gene set enrichment analysis of transcription factor targets from differentially expressed genes in scRNA-Seq data (**Fig. 2.5D**), for cells classified as erythroid progenitors, comparing treatment with Epo + IL-17A to Epo. Line indicates a significance threshold with 5% BH-corrected p-values, calculated using Fisher’s exact test. Colors distinguish spleen (SP) and bone marrow (BM) results. **F** Volcano plots showing differentially-expressed genes (grey points), and differentially-expressed pSTAT3 target genes (yellow points) in erythroid progenitors from spleen and bone marrow in Epo + IL-17A vs Epo conditions.

To validate the scRNA-seq results, we quantified changes in gene expression by qPCR in sorted CFU-E from bone marrow and spleen (4 mice per treatment). Of the four Epo-responsive genes selected for analysis (*Podxl, Cda, Arhgdig*, and *Cpd*), three were significantly elevated in CFU-Es following Epo+IL-17A treatment compared with Epo alone (**Fig. 6C**). Additionally, we measured cell-surface Podxl by flow cytometry and found it was increased by the combined Epo+IL-17A treatment compared with Epo alone, in BFU-E, CFU-E and ProE (**Fig. 6D**).

These observations strongly suggest that IL-17A acts by amplifying the Epo transcriptional response. To further test this, we carried out a gene-set enrichment analysis for canonical and non-canonical Epo^69^ and IL-17A^70^ signaling pathway response genes in the scRNA-Seq data (Table S4). The analysis revealed that, along with Stat3 and NFκB target genes, targets of Stat5, the canonical EpoR-activated transcription factor, are differentially enriched in erythroid progenitors from Epo + IL-17A-treated mice as compared to those treated with Epo alone (**Fig. 6E,F**). These results are consistent with our earlier work, which showed that Epo+IL-17A synergized in the phosphorylation of both Stat3 and Stat5 in erythroid progenitors and precursors^13^.

Collectively, these analyses show that IL-17A amplifies Epo-dependent gene expression, consistent with IL-17A-mediated sensitization of the response of erythroid progenitors to Epo (Model 2, **Fig. 4A**). The weak transcriptional responses in erythroid progenitors to IL-17A treatment alone are inconsistent with direct, Epo-independent action of IL-17A in these cells (as predicted by Model 1).

### The accelerated response to hypoxia by IL-17A is associated with faster cycling

To date, there is little direct data on the cell cycle in CFU-E and BFU-E *in vivo* during the response to Epo or hypoxic stress. Here we used our scRNA-seq data to infer progenitors’ cell cycle phase following each treatment (see Methods). We found increased number of erythroid progenitors in G2 or M following the combined treatment with Epo+IL-17A, compared with all other treatments (**Fig. 7A**). As a complementary approach, we calculated an activity score for genes canonically expressed in G2 or M. The G2/M score increased in erythroid progenitors in response to the combined Epo+IL-17A treatment compared with Epo alone, in both bone marrow and spleen (**Fig. 7B**).

**Figure 7.**
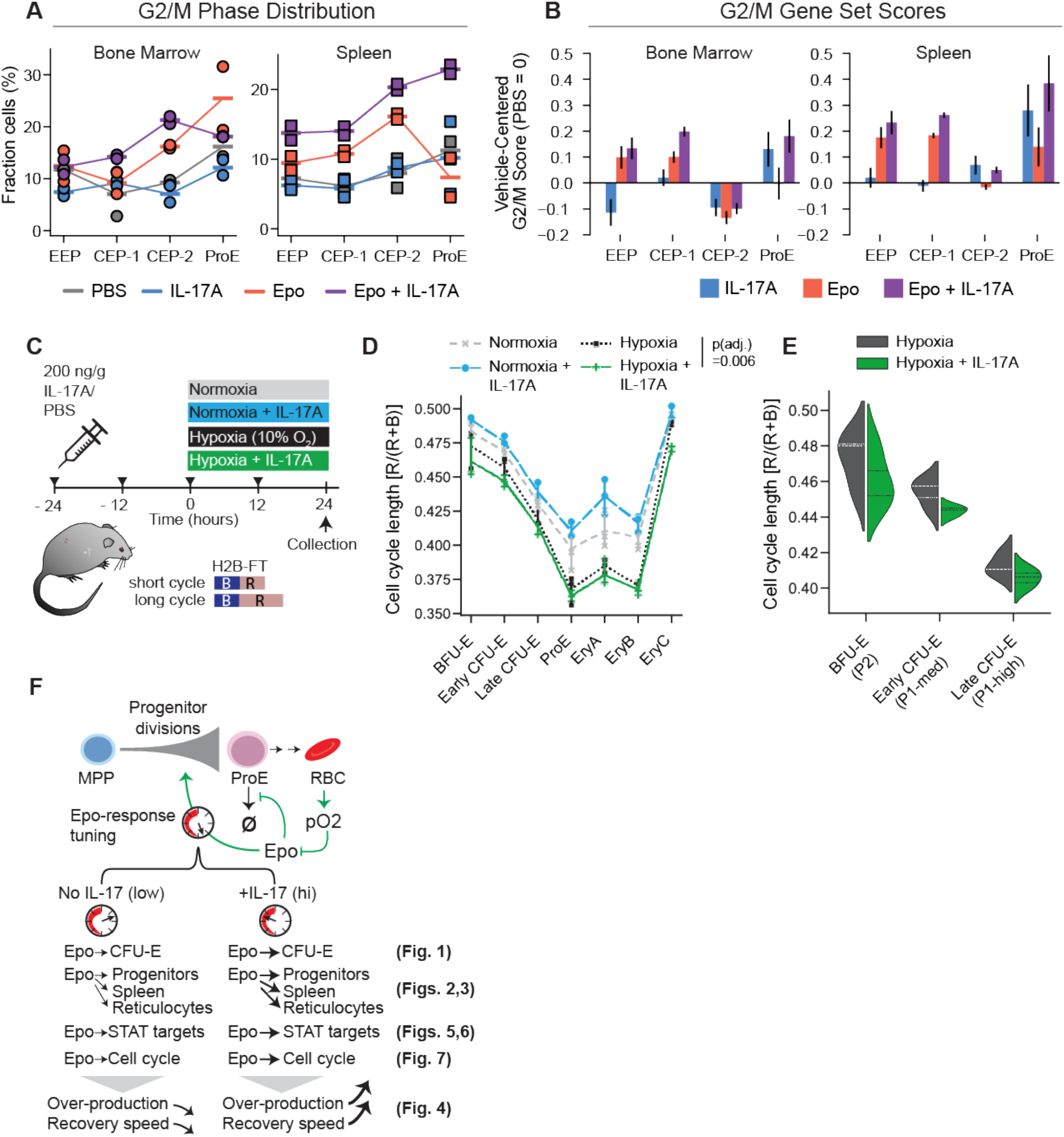
IL-17A enhances hypoxia-induced cell cycle shortening in erythroid progenitors. **A,B** The fraction of cells in G2/M phase (A), and expression of canonical G2/M-enriched genes (B), are elevated following joint Epo+IL-17A treatment as compared to treatment with IL-17A or Epo alone. Fractions and gene set scores are inferred from scRNA-Seq profiles in each of erythroid pro-genitor states (EEP, CEP-1, CEP-2, ProE) in bone marrow and spleen. **C** Experimental design for evaluating changes in erythroid progenitor cell cycle duration in response to hypoxia with or without IL-17A pre-treatment, using mice expressing the H2B-FT transgene, which undergoes blue(B)-to-red(R) maturation. The ratio of red to total (red and blue) H2B-FT fluorescence indicates cell cycle length, with a shorter cell cycle corresponding to a lower R/(R+B) ratio. **D** Median cell cycle length in the indicated bone marrow erythroid subsets was measured in n=3 or 4 mice for each treatment. P_adj.-_value is Wilcoxon signed-rank test (BH corrected), comparing erythroid subsets in hypoxia injected with either IL-17A or vehicle. **E** Distribution of cell cycle length in early erythroid subsets in hypoxia, injected with either IL-17A or vehicle. Data **Figure 2.7 (Continued)** pooled from 3 mice for each treatment. An average of 1300 cells (range: 328 to 5200) per erythroid subset per treatment were analyzed. **F** A model of homeostatic tuning by IL-17A. In baseline conditions, Epo regulates late precursor survival and division, while early progenitors are not dependent on Epo and undergo some Epo-dependent cell cycle shortening. In the presence of IL-17A, early progenitors become sensitized to Epo. They cycle more rapidly in hypoxia, show evidence of faster response, at the cost of further progenitor over-production.

These results indicated possible increased cycling in erythroid progenitors in response to the combined Epo+IL-17A treatment compared with Epo alone, and no corresponding response to IL-17A treatment alone. To investigate this further, we examined how IL-17A impacts erythroid progenitors cycling during hypoxic stress and in normoxia. We used mice transgenic for a live cell reporter of cell cycle duration, a chimeric histone H2B fused to a fluorescent timer protein (H2B-FT, **Fig. 7C**)^71^. H2B-FT fluoresces blue when first synthesized but matures over ∼1.5 hours into a red fluorescent protein. The ratio of red to total fluorescence is proportional to cell-cycle duration^71^. We treated H2B-FT transgenic mice with either IL-17A or PBS for 48 hours. After the initial 24 hours, half the mice from each group were transferred from normoxia to hypoxia (10% oxygen, n=3 or 4 mice per group, **Fig. 7C-E**). Using flow cytometry, we found that IL-17A by itself appeared to slightly increase cell cycle duration in erythroid progenitors, while hypoxia resulted in a clear shortening of the cycle in all erythroid progenitor and precursor subsets. The combined treatment of IL-17A and hypoxia resulted in a further cell cycle shortening, most pronounced in BFU-E and CFU-E (**Fig. 7D**, E; p_adj_= 0.006, Wilcoxon signed-rank test comparing all erythroid subsets in hypoxia treatment with the combined hypoxia + IL-17A treatment). We conclude that hypoxia promotes faster cycling in erythroid progenitors, an effect that is further enhanced by the addition of IL-17A. These findings are consistent with the hypothesis that IL-17A tunes the Epo feedback control by sensitizing erythroid progenitors to Epo, underlying the synergistic effect of Epo and IL-17A in progenitor proliferation and the stress response.

## Discussion

Our work establishes the IL-17A cytokine as an erythropoiesis stimulating agent that accelerates and augments the erythropoietic response to Epo and to hypoxic stress. Acting via the IL17RA/IL17RC heteromeric receptor, IL-17A sensitizes progenitors to Epo proliferative signals, thereby tuning the Epo-mediated feedback loop to stimulate erythropoiesis (**Fig. 7F**). IL-17A and Epo act synergistically in early erythroid progenitors, as seen by evaluating population changes through flow cytometry, transcriptional responses through scRNA-Seq and cell cycle speeds through a transgenic reporter. IL-17A has minimal effect on these progenitors when administered alone and it is not required for constitutive erythropoietic control. And yet in hypoxia or following Epo injection it has strong effects.

The interaction of IL-17A with Epo may shed light on some properties of tissue progenitor feedback circuits. The basic requirement for a homeostatic cell circuit is that it can match cell supply to tissue demand, but this requirement can be achieved with different reaction speeds, baseline fluctuations, and costs incurred by maintaining a proliferative compartment. In erythropoiesis, a cost-performance trade-off is evident as the system constitutively over-produces its own progenitors. Erythropoiesis begins with the long-lived self-renewing HSCs and MPPs, which differentiate through transit-amplifying erythroid progenitors into terminally-differentiating ProE, erythroblasts, and RBCs. This differentiation process lasts ∼7 days in mice and over 10 cell divisions ^23,72^, introducing a delay in the response to stress. This delay explains the continuous over-production of ProE ^20^, most of which undergo apoptosis in the absence of hypoxic stress. A dynamical model of erythropoiesis indeed confirms that continuous over-production accelerates response to stress. Accordingly, the action of IL-17A is consistent with pre-emptively further upregulating overproduction of ProE, thus accelerating the speed of response to hypoxia, should it occur, reducing both the depth of incurred hypoxemia and the time to recovery. This scenario may be relevant to infectious pneumonia, where IL-17A is an early marker of severe disease ^52-54^ and where hypoxemia, when it develops, does so progressively and is highly correlated with mortality ^73-75^. We indeed found that priming mice with IL-17A for 24 hours accelerated their erythropoietic response to subsequent hypoxia, an experimental model that may reflect the role of IL-17A during infectious pneumonia.

How does IL-17A tune the trade-off between ProE over-production and the speed of response to stress? By modeling, we showed that tuning components of the core Epo feedback can accelerate hypoxic responses through two effective mechanisms. IL-17A might act on the circuit to increase erythroid progenitor over-production independently of Epo (Model 1, **Fig. 4A**), or it might act to increase the sensitivity or response of erythroid progenitors to Epo (Model 2). A clear difference in the predicted behavior of these two models is that Model 2, tuning through Epo sensitization, accelerates the response to stress with a far lower burden of ProE overproduction than Model 1. Our experimental observations provided multiple lines of evidence that were strongly in favor of IL-17A acting by this efficient mechanism to handle the trade-off between ProE overproduction and a fast stress response. First, the response of the early progenitor pools (BFU-E, early and late CFU-E) to the combined treatment with both Epo + IL-17A was super-multiplicative, exceeding the product of response to each cytokine alone, as predicted by Model 2 but not by Model 1 (**Fig. 4E-G**). Second, in-depth single cell transcriptomic analysis showed that IL-17A amplifies the magnitude by which Epo and Stat5 gene targets are induced in early progenitors, while there was little transcriptional response to IL-17A alone in these cells. And third, we found that Epo shortened early progenitors’ cell cycles, and that these were further shortened by the combined Epo + IL-17A treatment, while IL-17A by itself did not shorten (and even somewhat lengthened) these cycles. The full molecular mechanism by which IL-17A sensitizes erythroid progenitors to Epo will require further elucidation, but is at least in part likely to be the result of synergistic activation of Stat3 and Stat5 by the combined action of Epo and IL-17A^13^.

There are several generalizations that might be made from this work. Tuning of over-production is used to accelerate responses in more general biological contexts. For example, when *E. coli* bacteria grow with abundant nutrients, they resort to ‘overflow’ metabolism that is inefficient but accelerates their growth rate ^76,77^. Our work shows that an overflow-like mechanism operates in the regulation of complex cellular circuits in higher organisms. The mode of action found here for IL-17A— sensitizing cells to Epo—explains how a pleiotropic factor like IL-17A may act with specificity. This specificity of action may allow IL-17A to be used in therapy in anemia. Beyond erythropoiesis, these findings show that IL-17A accelerates response in one lineage with minimal effects on others, and yet the *Il17ra* receptor is broadly expressed and is documented to amplify signals of other tissue-specific factors ^60,78,79^. Our work thus offers a template for understanding these broader functions of IL-17 and suggests a framework for yet other cytokine interactions as tuning performance goals.

## Acknowledgements

We thank Jun Huh and Eunha Kim at Harvard Medical School for providing the IL-17RA germline knockout mice. We thank the Harvard Medical School Immunology Flow Cytometry Core Facility for their guidance on panel design and spectral analyzer training. We thank the UMass Chan Flow Cytometry Core for their expertise and NIH S10OD028576 for the purchase of the BDFACSFusion Cell Sorter.

## Declaration of Interests

AMK is a cofounder of Somite Therapeutics.

## Funding

This work was supported by the National Institutes of Health [R01HL141402 to MS and AMK; R01DK120639 to MS] and by a Harvard QFASTR grant. AMK acknowledges support of a Mallinckrodt Foundation Scholarship. QCW received support from the National Science Foundation Graduate Research Fellowship Program [DGE2140743 and DGE1745303] and the Herchel Smith Graduate Fellowship

## Materials and Methods

### In vivo animal studies

Mice treated with IL-17 and Epo in vivo were either BALB/cJ, Il17ra^f/f^/Vav-iCre (obtained by crossing Il17ra^f/f^ (B6.Cg-Il17ra^tm2.1Koll^/J) with Vav-iCre (B6.Cg-Il17ra^tm2.1Koll^/J)) or Vav-iCre. Mice treated with IL-17 and hypoxia were either BALB/cJ, or mice transgenic for H2B-FT (B6;129-Gt(ROSA)26Sor^tm1(rtTA*M2)Jae^ Hprt1^tm2(tetO-mediumFT*)Sguo^/Mmjax). Germline Il17ra deletion mice were provided by Jun Huh at Harvard Medical School (obtained by crossing exon 4 to 7 floxed Il17ra^f/f^ strain with Ubc-Cre-ERT2 (B6.Cg-Ndor1^Tg(UBC-cre/ERT2)1Ejb^/1J, and outcrossed with B6 mice to remove Ubc-Cre) as previously described ^1,2^. All mice were 8 to 12 weeks old at the time of experiments. Littermates of the same age and sex were randomly assigned to treatment groups. Both male and female mice were used in experiments. The experimental protocols were approved by the University of Massachusetts Chan Medical School Institutional Animal Care and Use Committee (IACUC) for the Socolovsky laboratory or by the Harvard Medical School IACUC for the Klein laboratory.

### Primary cultures

Colony-formation assays were done with bone-marrow harvested from 8-to 12-week-old male or female BALB/cJ bone marrow.

### CFU-E colony formation assays

Adult mouse bone-marrow cells were plated at 500,000 cells/ml in methylcellulose (made in-house, containing Iscove’s Modified Dulbecco’s Medium (IMDM), 6% Bovine serum albumin (BSA), 5% fetal calf serum (FCS), 1mg/ml human transferrin) with added Epo and IL-17 cytokines at the indicated concentrations. Colonies were counted at 72 hours after staining with diaminobenzidine to visualize hemoglobinized cells.

### Cytokine treatment in vivo

Mice were injected with cytokines subcutaneously, in a total volume of 4 microliters/g body weight. Recombinant IL-17A (made in Chinese Hamster Ovary (CHO) cells) was freshly resuspended in phosphate buffered saline (PBS) up to 12 hours prior to use and injected at 200 ng/g body weight every 12 hours. Epo (Procrit) was injected at 0.25 IU/g every 24 hours. Vehicle-treated mice were injected with an equal volume of PBS.

### Tissue Harvesting

Femur, tibiae, and spleen were harvested following euthanasia and submerged in SB5 buffer (phosphate-buffered saline (PBS) supplemented with 0.2% (w/v) bovine serum albumin (BSA), 0.08% (w/v) glucose, and 5 mM of Ethylenediaminetetraacetic acid (EDTA)). Tissue harvest methods are described in detail in Swaminathan et al.^3^. In brief, for bone marrow samples, the ends of the bones were cut and flushed with SB5 using a 26-gauge needle and 5 mL syringe. Harvested bone marrow was filtered through a 100 µM cell strainer. For spleen samples, spleens were placed onto a staining buffer-wetted 100 µM cell strainer and macerated through using the rubber side of a syringe plunger. The samples were washed with more SB5 and pelleted at 2000rpm for 10 minutes. Bone marrow and spleen samples were then processed into single-cell suspensions by adding 2 mL of SB5 and triturating cell pellet using a P1000 pipette and a large orifice pipet tip.

### Hypoxia model of erythropoietic stress

Mice were placed for up to 48 hours in a hypoxic environment using the BioSpherix A chamber (BioSpherix). Hypoxia was achieved by displacing oxygen with nitrogen at normal atmospheric pressure. Temperature, humidity, and carbon dioxide readings were monitored.

### Flow cytometry

All antibody panels included either Fc block or rabbit IgG, and all cells were post-labeled with DAPI to dead cells. Reticulocytes were measured by flow-cytometric analysis of blood labeled with the DNA stain Draq5 and with CD71 antibody. Erythroid precursors (ProE, EryA/B/C) were identified in spleen and bone marrow samples that were labeled with lineage markers (Gr1, Mac1, CD4, CD8a, CD19, F4/80), CD71 and Ter119 antibodies^4^. Early hematopoietic and erythroid progenitors (BFU-E/P2, early CFU-E/P1-medium, late CFU-E/P1-high, basophil/mast cell progenitors/P3, megakaryocytic progenitors /P4, EBMP/P5) were identified by labeling with antibodies directed at Kit, lineage markers, Ter119, CD71, CD55, CD49f, CD105, CD150, CD41 (the ‘10 color panel’) ^2,3^. The Podxl antibody was added to the 10-color stain panel in the indicated experiments. Early hematopoietic progenitors CMP, MEP and GMP were identified using antibodies directed at Kit, Sca1, lineage markers, CD34 CD16/32. Non-erythroid hematopoietic cells (‘lympho-myeloid’ subsets) were identified using antibodies directed at CD19, CD4, CD8a, Mac1 and Gr1. All antibody labeling was done on freshly harvested cels at 4°C. Cells were analyzed on a Cytek Aurora cytometer with 5 Lasers. Data was analyzed using FlowJo 10.10.0.

### Flow sorting

For isolating BFU-E, CFU-E, or Kit^+^: harvested bone-marrow and spleen cells were enriched for Kit^+^ early progenitors using MojoSort Streptavidin Nano-bead bound to biotinylated anti-CD117 antibody. Enriched cells were then labeled with the ‘10 color panel’^2,3^ antibodies and sorted on a BD FACSFusion with 5 Lasers. For isolating ProE, harvested spleen and bone-marrow were labeled with CD71, Ter119 and lineage markers and sorted or on a BD FACSFusion.

### qPCR assay for deletion efficiency of Il17ra

Genomic DNA was extracted from whole bone marrow and spleen, or from flow-sorted ProE and Kit+ cells. For each genomic DNA sample, qPCR was performed using unique primers to exons 2 or exon 3 within the ‘floxed’ regions, and to either exon 7 or exon 5 outside the ‘floxed’ region. Deletion efficiency was calculated from DDCT between the exons that are external and internal to the ‘floxed’ region.

### IL-17A serum measurements

Murine IL-17A from blood serum was measured using the Mouse IL-17A ProQuantum Immunoassay Kit following the manufacturer’s protocol. The CFX96 Touch Real-Time PCR Detection System was used for the readout.

### Dynamical Systems Modeling of Tuning in Erythropoietic Differentiation

Definition of the dynamical systems model shown in **Figs. 4**, S6 is provided in the file Supplementary Text 1, including details on generation of all modeling figure panels. Numerical simulations were performed as described in this supplement using Python (v3.10.13) with SciPy (v1.11.4), with code available at github.com/AllonKleinLab/Wu2024.

### Singe cell RNA sequencing (scRNAseq) of bone marrow and spleen cells

#### CD117 positive cell enrichment

Harvested tissue samples were first enriched for CD117+ cells. Bone marrow and spleen cell samples were washed using Easy Sep buffer (PBS, 2% fetal bovine serum (FBS), and 1 mM EDTA) and enriched for CD117 expressing cells using EasySep™ Mouse CD117 Positive Selection Kit. The enrichment was performed largely following the manufacturer’s protocol; however, all room temperature steps were instead performed at 4°C in a walk-in cold room.

#### Live cell enrichment using density centrifiugation

Following CD117 enrichment, dead cell and debris were removed using gradient centrifugation in OptiPrep. Solutions of 40%, 18%, 12%, and 5% OptiPrep were prepared using weight by volume (w/v) and pre-chilled. Cells were resuspended in 0.5 mL PBS and mixed with 1 mL of 40% OptiPrep in a 5 mL FACS tube. 18%, 12%, and 5% OptiPrep were sequentially and carefully overlay on-top of the cell sample. The samples were then spun at 800g for 15 minutes with the centrifuge break off to prevent disruption of gradient. The live cell layer, which formed between the 40% and 18% interface, was collected, and washed using cold PBS.

#### MULTIseq labeling ofi cell samples

To reduce technical variability from sample processing, we performed MULTIseq labelling to multiplex samples for 10X Chromium-based encapsulation. Samples were labeled with MULTI-Seq Lipid-Modified Oligos and unique sample barcode oligos following the MULTI-Seq protocol^5^ (see Table S4 for MULTI-Seq barcodes per sample and Table S5 worksheet “Primers for Hashtags” for primer sequences used). Antibody-based hashing methods were tested for these cell samples but ultimately not used due to poor labeling of reticulocytes which lack of the hashing epitope.

#### Single cell encapsulation and library prep

Following MULTI-seq sample labeling, samples were pooled in equal ratios and cells were resuspended in PBS and 0.04% BSA at 1000 cells/uL prior to cell encapsulation. To help with demultiplexing, single injection condition pools were run individually, and an all-sample pool was run in another, for a total of five unique pools and a target total cell number of 48,000 loaded separately onto a 10X Genomics Chromium Next GEM Chip G. Samples were encapsulated using a Chromium X Instrument based on manufacturer’s instructions.

The scRNA-seq gene expression libraries were processed using the standard procedures following the Chromium Single Cell 3’ Reagents Kits User Guide (v3.1 Chemistry). The MULTI-seq barcode libraries were processed following the MULTI-seq library preparation protocol ^5^. Library QC was performed using Agilent 2100 Bioanalyzer, Agilent Tape station, and KAPA Library Quantification. Libraries were sequenced on the Illumina NovaSeq 6000 using an S4 200 cycle flow cell and SP 100 cycle flow cell with a target read depth of 25,000 reads per cell for the gene expression and 5,000 reads per cell for the sample barcodes.

### scRNAseq Data Analysis

#### Computing environment

Unless stated otherwise, all analyses were carried out in Python (v 3.7.12), using pandas (v1.3.5) for data manipulation, numpy (version 1.21.6) and scipy (version 1.7.3) for numerical computations, and scanpy (version 1.9.1) for handling single-cell data. These analyses were performed on Harvard Medical School’s O2 high-performance computing cluster. Computational efficiency was enhanced by parallelizing computations across cell states using the multi-processing.

#### Data Preprocessing

The sequencing libraries were demultiplexed using Bcl2fastq (Illumina) and concatenated across multiple sequencing runs. The demultiplexed Fastq files were processed using Cellranger (v6.1.2) Gene Expression Pipeline. Alignments were performed using M. musculus Ensembl release 84 mm10 v1.2.0 cDNA reference. The MULTI-seq library Fastq files were processed using CITE-seq-Count tool (v1.4.5)^6^ to determine sample identity of pooled cells. Sample barcode identity assignments to cells were performed using a custom python-based adaption inspired by the deMULTIplex R package^5^.

Gene expression data quality control was performed using the Scanpy pre-processing pipeline with the following criteria: sc.pp.filter_cells was used to include cells with at least 500 unique molecular identifies (UMIs) and 200 unique genes expressed. Cells with more than 5% transcriptome contributed by Gm genes (Gm26917, Gm42418, Gm25580, Gm24139, Gm24146) or by mitochondrial genes (beginning with mt-), representing likely stressed or dead cells, were excluded from downstream analysis. Doublets were identified using Scrublet^7^ and also filtered from further processing. Sc.pp.filter_genes was used to filter for genes that were expressed in at least 3 cells across all samples. The gene expression matrix was then globally scaled by normalizing gene expression measurements by the total UMIs per cell and multiplied by a scaling factor of 1e4 to obtain counts per ten thousand (CP10K). Gene expression values were then transformed to log(1+CP10K).

#### Transfier learning and UMAP visualization

For data graph embedding and visualization, we first projected the transformed data into a linear subspace learnt by carrying out principal component analysis (PCA) on an scRNA-seq dataset from a well-characterized murine hematopoiesis reference (GEO accession: GSE89754) ^2^. To carry out PCA on the reference data set, we processed the raw counts following the same steps as described above and used highly variable genes (selected using scanpy’s sc.pp.highly_variable_genes with default parameters) to perform dimensionality reduction by PCA (for N=50 components). Using the PCA loadings of the reference dataset, we then projected the new dataset collecting in this study, from all conditions, onto the reduced principal component space. A kNN graph was constructed for the pooled samples using k=10 neighbors and then visualized using UMAP ^8^.

#### Cell State Annotations

Cell annotations were assigned using a combination of query-based transfer learning and manual curation. Published scRNA-seq data sets containing bone marrow and splenic cells were curated and used as reference data sets^2,9,10^ (GEO accessions: GSE89754, GSE132042, GSE132901, GSE261601). Each reference dataset used for cell annotation was pre-processed individually in the method described above for the reference data set used for visualization.

The new data set was projected into a PCA subspace constructed for each of these reference data sets, following the same steps as in the “Transfer learning and UMAP visualization” subsection above. This process resulted in four independent rounds of classification. In each case, a k-nearest neighbors (kNN) classifier (k = 5), using majority voting^2^, within each neighborhood, was applied in each round to assign annotation labels from the corresponding reference dataset. Thus, each cell received four separate labels, one from each round of classification. After obtaining labels from multiple reference data sets, labels were confirmed by manual inspection of cell type-specific marker genes (Table S4). In cases where annotations between the four reference sets disagreed, Leiden clustering (using sc.tl.leiden with resolution = 2) was performed to sub-cluster populations with ambiguous labels. Marker genes were identified for each ambiguous sub-cluster using sc.tl.rank_gene_groups, and a literature search was conducted to confirm the correct annotation based on marker genes. Condition Dependent Embedding Density Mapping

To identify regions of enrichment or depletion of cell states in our data, we calculated embedding densities for each treatment (**Fig. 5C**). A k-nearest neighbors (kNN) graph (k = 200) was constructed using the principal component (PC) space defined in the dimensionality reduction section. The adjacency matrix from this graph was used to estimate a probability density function (PDF) for each condition.

Let *A*_*i,j*_ = n, n ∈ [0, *k* − 1] represent the number of neighbors for cell i that belong to treatment j in a neighborhood of size k. To control for sampling biases, we applied a two-step normalization process. We first normalized A column-wise to obtain,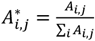. This adjusts for differences in the total number of cells per treatment, representing the neighborhood composition as a normalized probability. Next, we performed row normalization to convert the treatment-dependent neighborhood compositions into probability densities for each cell, ensuring that the contributions from all treatment within a cell’s neighborhood sum to 1. Each value, 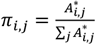, represents the probability that a neighbor of cell i comes from treatment *j*.

To quantify treatment-specific enrichment relative to the control (PBS or Epo), we computed the *log*_2_ fold change between the probability densities of the perturbed treatment and the control:

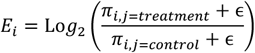

Where ∈ = 10^−4^ is a pseudocount to avoid divisions by zero.

The calculated enrichment densities per cell, *E*_*i*_, were plotted on the UMAP embedding to visualize the global differences across treatments. Differential Gene Expression Analysis

Differential gene expression analysis (**Fig. 5D**) was conducted using Scanpy’s sc.tl.rank_gene_groups function, using two-sided Wilcoxon rank-sum test on log-normalized gene expression values. Each biological replicate was paired with a matching control replicate. The analysis was performed on a cell state-by-cell state basis across conditions and across organs. To account for biological replicates, the raw p-values obtained from Wilcoxon rank-sum test were combined between replicates using Fisher’s Method. A false discovery rate (FDR) was calculated using the combined p-values using the Benjamini-Hochberg procedure. A gene was considered differentially expressed if it had an absolute log2(FC) > 0.25 between the cytokine treatment-comparisons in both biological replicates and whose was FDR < 0.05. Mitochondrial genes, unlabeled genes (genes starting with Gm, AY, AC), and genes whose detection is high in empty droplets (indicative of ambient RNA) were excluded.

#### Generalized Linear Model fior Formal Assessment ofi Cytokine and Organ Contributions

To determine the overlap of IL-17A gene expression response with Epo response while controlling for biological differences between organs (spleen vs bone marrow), we performed generalized linear modeling (GLM) on the scRNAseq data. The GLM was fit separately for each cell annotation: early erythroid progenitors (EEP), and two subsets of committed erythroid progenitors (CEP-1 and CEP-2). Genes expressed in fewer than 10 cells were excluded. The GLM was applied to the log(1+CP10K) gene expression values.

A design matrix was constructed using binary indicator variables to encode metadata variables for treatment conditions and organ origin. For treatments, cells treated with IL-17 were assigned a value of 1 for the IL17 variable and 0 otherwise, similarly for EPO with the EPO variable. Organ origin was encoded with 1 representing spleen and 0 representing bone marrow (reference organ) in the Organ variable. The model has the form:

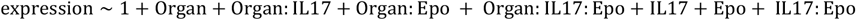

with a gaussian noise. To identify genes whose expression is regulated by specific factors and interactions, we used a Likelihood Ratio Test comparing to nested (‘null’) models defined in **Fig S7B**. For each nested model we calculated p-values for each gene from a chi-squared test on the LRT test statistic:

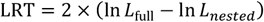

where In *L*_fuII_ and In *L*_*nested*_ are the log-likelihoods of the full and nested models, respectively. The p-values were then adjusted to control for the false discovery rate (FDR) using the Benjamini-Hochberg procedure. Correction was performed separately for each cell state. All statistical analyses were conducted using Python (version 3.9.19), using pandas (version 1.4.3) for data manipulation, numpy (version 1.23.0) and scipy (version 1.11.0) for numerical computations, and scanpy (version 1.9.8) for handling single-cell data. The statsmodels package (version 0.14.3) was used for fitting generalized linear models (GLMs), using sub-routine GLM.

#### Pseudotime analysis

Erythroid pseudotime analysis (**Figs. 6A,B**) was performed on a subset of the data that only includes cells annotated as: MPP, EBMP, EEP, CEP-1, CEP-2, ProE, BasoE, PolyE, OrthoE, and Retic. Pseudotime was assigned to each cell using Scanpy’s scanpy.tl.dpt, with a randomly-selected MPP was used as the root cell to define the start of the trajectory.

#### Epo Response and Cell Cycle Phase Gene Set Scoring

Gene sets were defined from previously published work and are given in Table S4 (worksheets “Gene Sets Used in Paper” and “Gene Set Gene List”). The Epo response gene set (**Figs. 6A,B**) was obtained from differentially genes from Epo stimulated progenitors in Tusi et al.^2^, and the cell cycle G2/M phase gene set (**Fig. 7B**) was selected from Whitfield et al. ^11^. For each of these two gene sets, scores (**Figs. 6A,B** and **7B**) were calculated using the scanpy.tl.score_genes function, providing gene expression after pre-processing to log(1+CP10K) (see ‘Data preprocessing’ above), and the following parameter values: ctrl_size=N where N is is equal to the size of the test gene set; gene_pool=all_non_set_genes, where all_non_set_genes is the set of all genes excluding the test gene set.

#### Gene Set Enrichment Analysis of Transcription Factor Targets

Gene set enrichment analysis (**Figs. 6E,F**) was carried out between cells from Epo + IL-17A treated mice and Epo treated mice. Only cells annotated as erythroid progenitors (EEP, CEP-1, CEP-2) were included in analysis. We used transcription factor target gene sets collected from literature search of CHIP-seq and RNA-seq publications, as well as MSigDB v5.1 curated sets (Hallmark and C2 chemical and genetic perturbations, C3 transcription factor targets)^12^ (see Table S4 for gene sets, worksheets “Gene Sets Used in Paper” and “Gene Set Gene List”).

Differentially-expressed genes were identified as described in section ‘Differential Gene Expression Analysis’ above. Prior to set-enrichment analysis, ribosomal genes, mitochondrial genes, and predicted genes (genes starting with “Gm”) were excluded from the input. Set enrichment was determined using Fisher’s exact test (one-tailed) and the resulting p-values were then adjusted using the Benjamini-Hochberg procedure across all gene sets to control for FDR.

#### Cell cycle phase assignment

For **Fig. 7A**, we further performed cell cycle phase assignments adapting methods described in Tirosh et al.^13^ Briefly, a score was calculated for each cell cycle phase (G1/S, S, G2, G2/M, M/G1) scores as described above, with gene sets given in Table S4 (worksheets “Gene Sets Used in Paper” and “Gene Set Gene List”). Each cell cycle phase score was separately mean-centered and standardized across all cells. Cells with phase scores across all phases below 0 were assigned to G0 as they cannot be clearly assigned to a cell cycle phase from this analysis. We assign the remaining cells to the phase which the phase score is the highest across all the scores.

RNA isolation and RT-qPCR of sorted CFU-Es (gate P1)

Due to the high level of ambient RNA present in FBS, sorted cells were washed twice in large volumes (4.5 mL) of ice-cold staining buffer to minimize RNA contamination. After washing, cells were pelleted by centrifugation, and the supernatant was removed. Cell pellets were flash-frozen and stored at -80°C until further analysis. Total RNA was extracted from frozen pellets using the RNeasy Micro Kit according to the manufacturer’s instructions. Whole-transcriptome pre-amplification was carried out prior to qPCR as follows: cDNA synthesis was performed using Maxima H Minus Reverse Transcriptase in the presence of a poly-T oligo and template-switching oligo (TSO). Whole-transcriptome cDNA amplification was performed using KAPA HiFi HotStart Ready Mix and cDNA amplification primers following the manufacturer’s protocol. Subsequently qPCR was carried out on the amplified cDNA using KAPA SYBR Fast at a final reaction volume of 10 µL, containing 2 µL of cDNA template, 5 µL of SYBR Green master mix, and 200 nM each of forward and reverse primers for the genes of interest. The primer sequences for each gene as well as the sample prep primers are listed in Table S5, worksheet “qPCR Primers for Epo Response”. Measurements were performed in either 96-well or 384-well qPCR plates. For 96-well plates, reactions were run on a CFX96 Touch Real-Time PCR Detection System. For 384-well plates, reactions were run on a QuantStudio 7 Flex Real-Time PCR System. All reactions were performed in technical triplicates.

### Quantification and statistical analysis

Statistical tests are indicated in the figure legend and text of each experiment. All tests that are part of multiple hypotheses were corrected for false discovery using the Benjamini Hochberg procedure as stated in text, with adjusted p-values given as p_adj_. Tests considered significant at a 5% false discovery rate (p_adj_ <0.05), or p<0.05 for isolated tests.

---

**Supplementary Figure 1.**
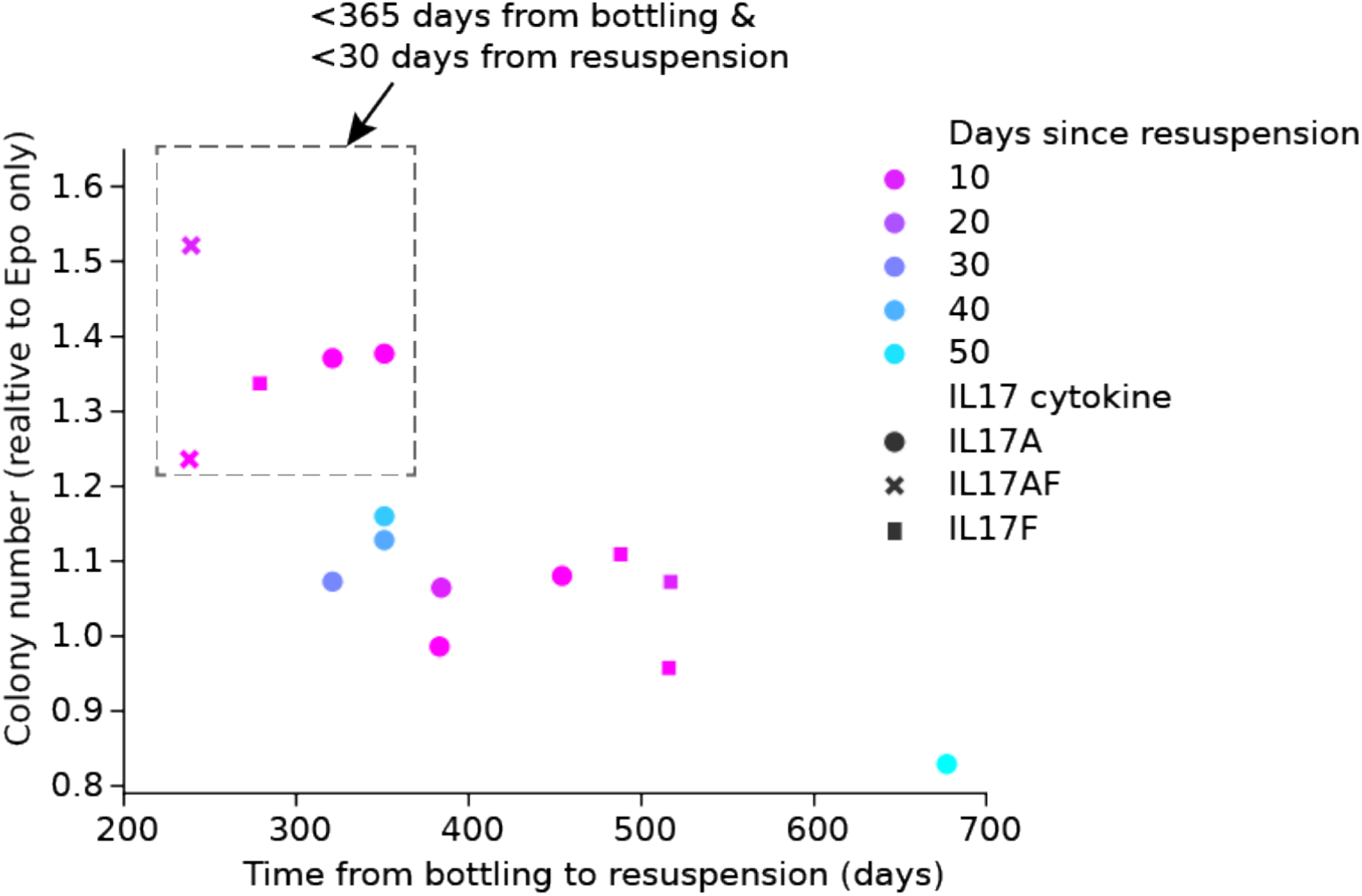
IL-17 cytokine activity in the CFU-e assay decays with the age of the IL-17 lyophilized preparation. CFU-E assays were carried out as in **Fig. 1E**. Lyophilized protein was purchased from R&D Systems. The time interval between bottling of the lyophilized protein by the manufacturer, and its resuspension prior to use, varied for different protein lots. Activity is lost if the lyophilized protein was bottled > 365 days prior to the day of resuspension. In addition, once the protein is resuspended, its activity declined rapidly after 30 days. All IL-17 protein ligands (both lyophilized and resuspended) were stored at -80°C.

**Supplementary Figure 2.**
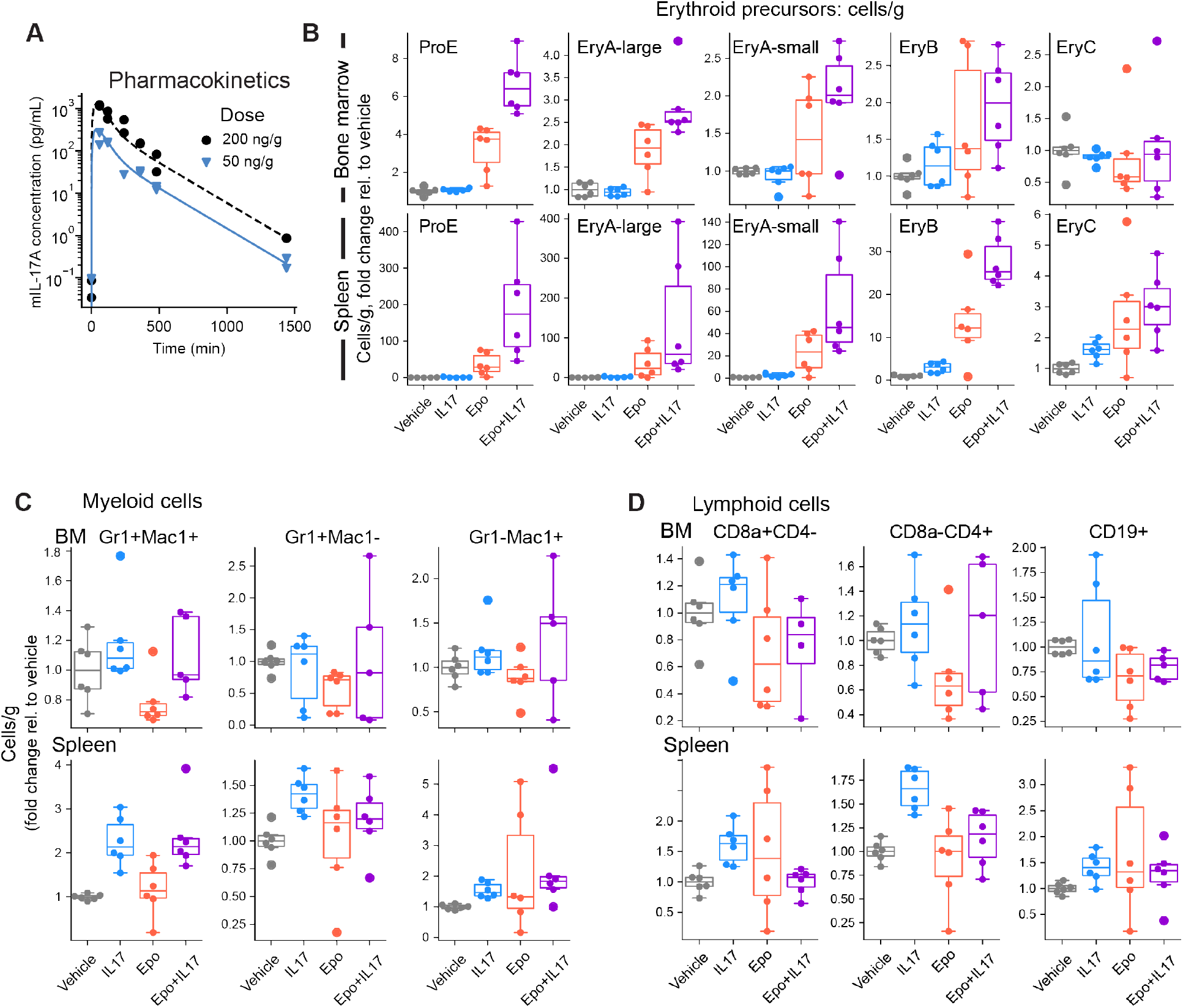
Flow cytometric analysis of erythroid precursors and lympho-myeloid cells in mice treated with Epo and IL-17A. **A** Pharmacokinetics of IL-17A *in vivo*. 2 mice per dose were injected with the indicated dose of IL-17A (50 ng/g and 200 ng/g). Blood was obtained from tail bleeds and IL-17A was measured in plasma in small volumes of plasma using immunoPCR. The figure is reproduced with permission from (Wu et al, 2024 in review). **B-D**, Data underlying **Fig. 2.2F**, broken out by mouse: **B**, Flow-cytometric analysis of erythroid precursors in terminal differentiation. Freshly harvested bone marrow and spleen cells were labeled with CD71 and Ter119 antibodies and immediately analyzed to identify ProE, EryA/B/C subsets. FACS gates as in^36^. Data points correspond to individual mice and box plots are defined as in **Fig. 2.2B. C, D** Flow-cytometric analysis of lymphoid and myeloid subsets in bone marrow and spleen.

**Supplementary Figure 3.**
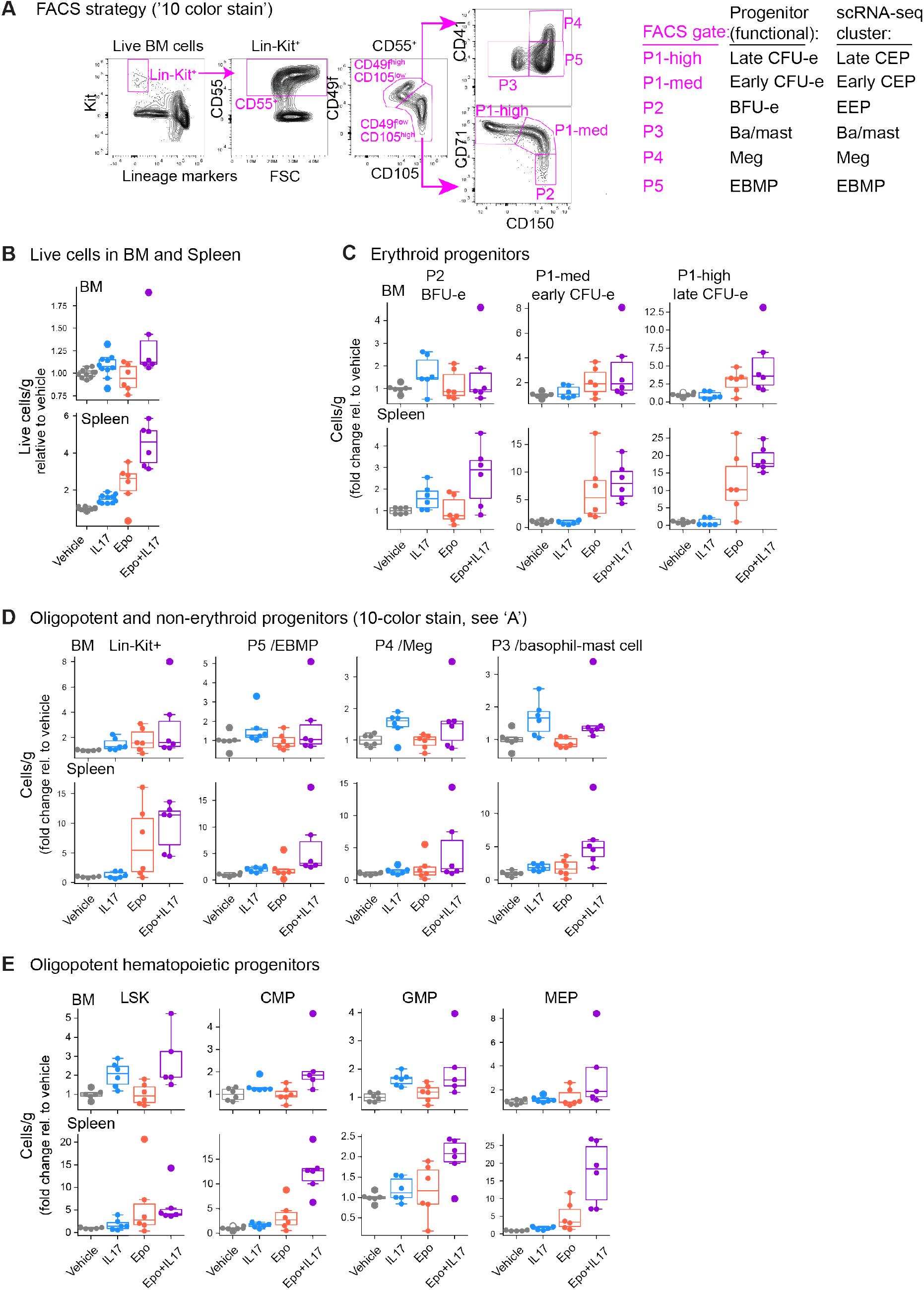
Flow cytometric analysis of early hematopoietic and erythroid progenitors in mice treated with Epo and IL-17A. **A** FACS gating strategy for identifying early hematopoietic and erythroid progenitors^13,37^ (the ‘10 color stain’). Corresponding FACS gates, functionally-defined progenitors, and single-cell RNA-seq (scRNA-seq) clusters are shown on the right. Correspondence between the three modalities of defining each progenitor type was previously shown in Tusi et al.^13^ **B-E** Data underlying **Fig. 2F**, broken out by mouse. **B-D**, Summary data of bone marrow (BM) and spleen cells labeled with the ‘10 color stain’ antibody panel. Data points correspond to individual mice and box plots are defined as in **Fig. 2B. E**, Summary data of BM and spleen cells labeled with CD34, CD16/32 antibodies. CMP, common myeloid progenitor; MEP, megakaryocytic/erythrocytic progenitor; GMP, granulocytic/monocytic progenitor.

**Supplementary Figure 4.**
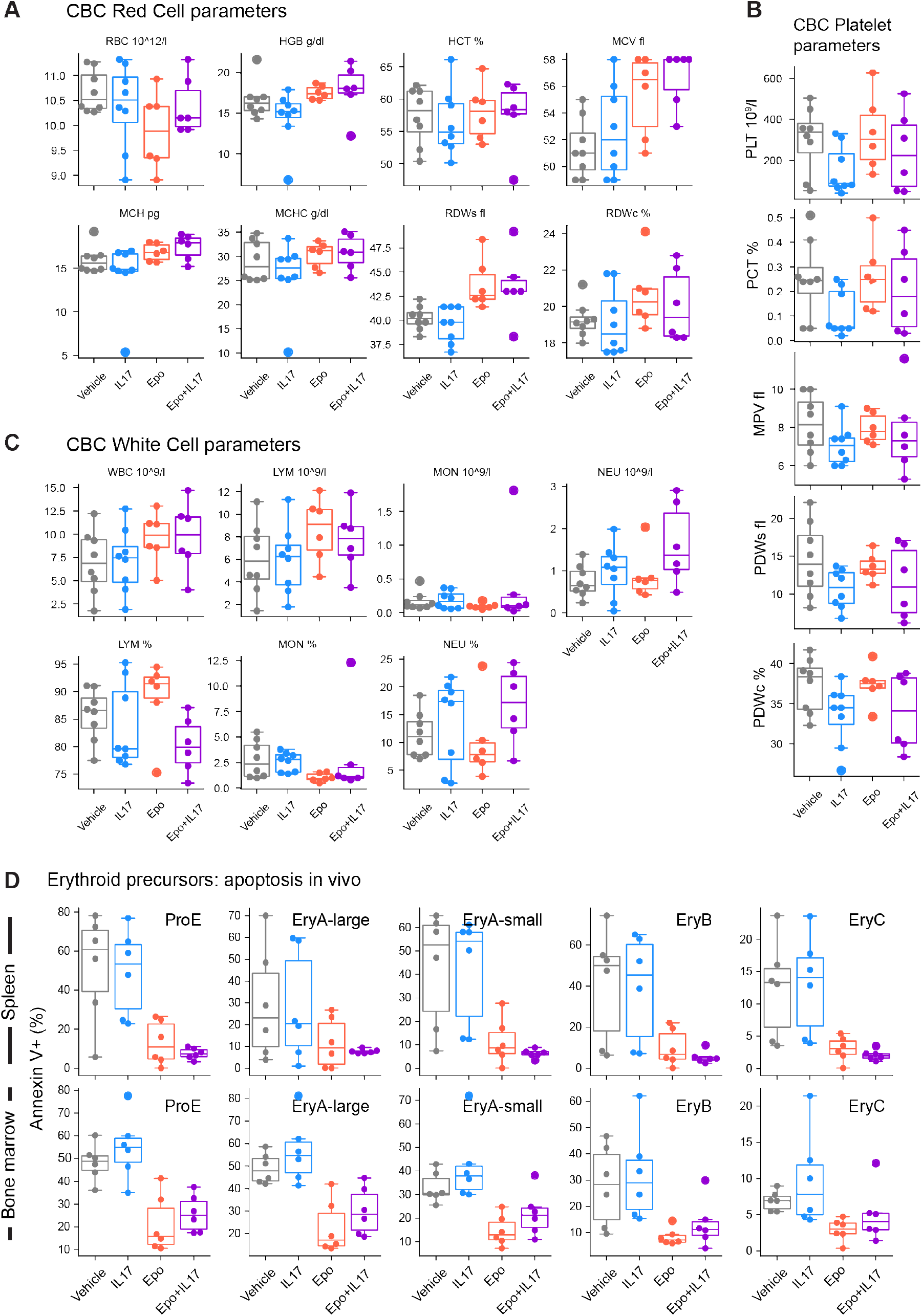
Complete blood count (CBC) and erythroblast survival measurements in mice treated with Epo and IL-17A. **A-C** CBC results from experiments described in **Fig. 2**. Blood was obtained by cardiac puncture immediately following culling. Data points correspond to individual mice and box plots are defined as in **Fig. 2B-D** Flow-cytometric analysis of apoptosis in erythroid precursors *in vivo*. Annexin V labeling of cells in each of the erythroid subsets shown in **Fig.S2B**.

**Supplementary Figure 5.**
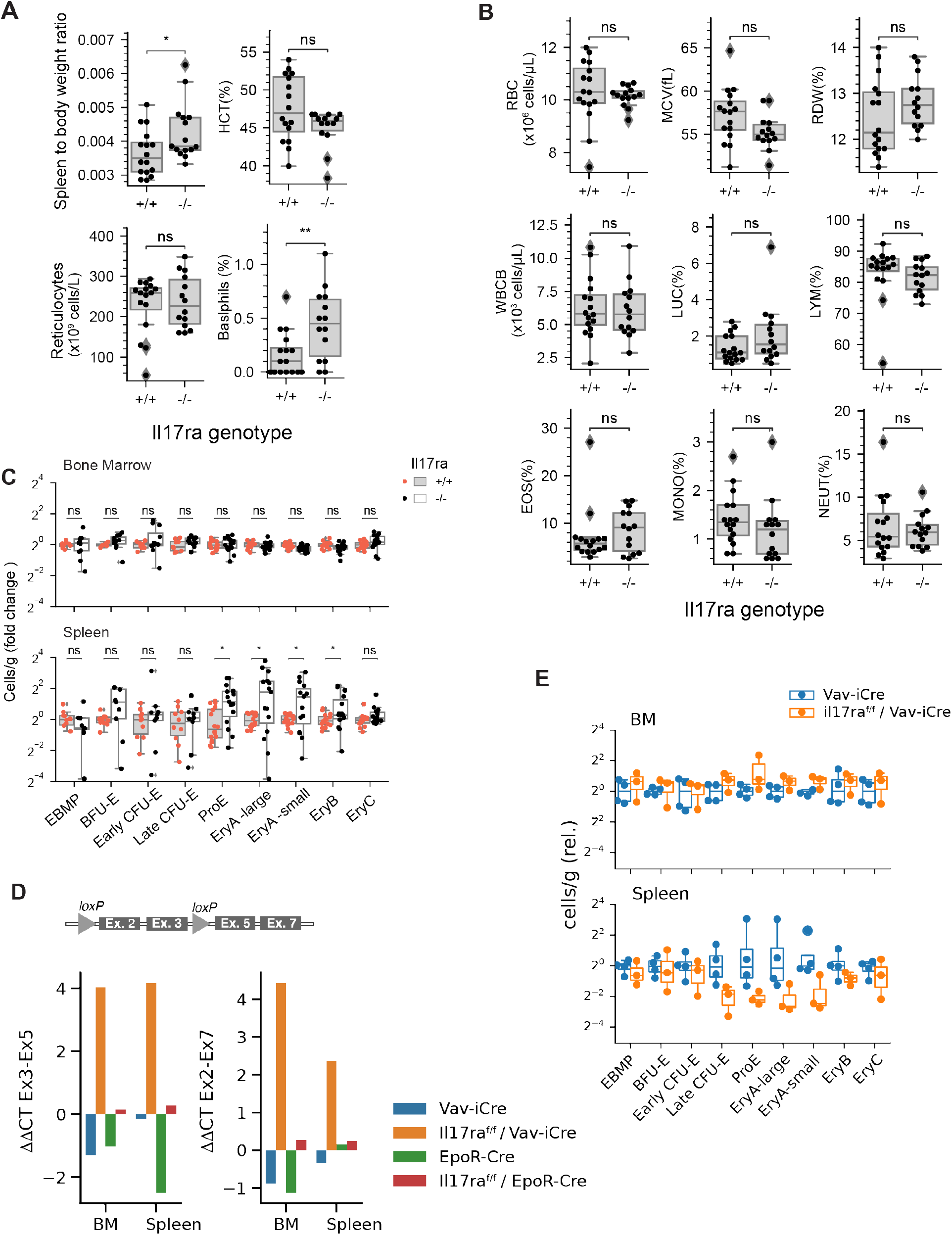
Germline and conditional deletions of *Il17ra*. **A**, **B** Blood and spleen parameters including complete blood count (CBC) analysis in germline deletion of ***Il17ra*** and in age- and sex-matched wild-type controls. P-values calculated using Wilcoxon rank sum test (* p < 0.05, ** p < 0.01, ns: not significant).**C** Bone marrow and splenic early erythroid progenitors and precursors in germline ***Il17ra*** knockout mice vs. age-and sex-matched controls, revealing extramedullary erythropoiesis. Flow cytometry analysis using CD71/Ter119 and ‘10 color stain’ panels (n=12-15 mice/condition). *p_adj_<0.05, ns: not significant (Wilcoxon rank sum test, Bonferroni correction). **D** qPCR analysis of *Il17ra* deletion assessed using whole bone marrow or whole spleen. Two alternative primer sets gave similar results: either primers to exons 3 and 5, or exons 2 and 7. For Vav-iCre -mediated deletion, deletion efficiency in whole tissue was less efficient than in sorted hematopoietic cells (either Kit^+^ or ProE), see **Figure 3**. EpoR-Cre -mediated deletion was poor. **E** Early erythroid progenitors and erythroid precursors in Vav-iCre and Il17ra^f/f^ / Vav-iCre, analyzed by flow cytometry using CD71/Ter119 and the ‘10 color stain’ panels (see **Fig. S3**).

**Supplementary Figure 6.**
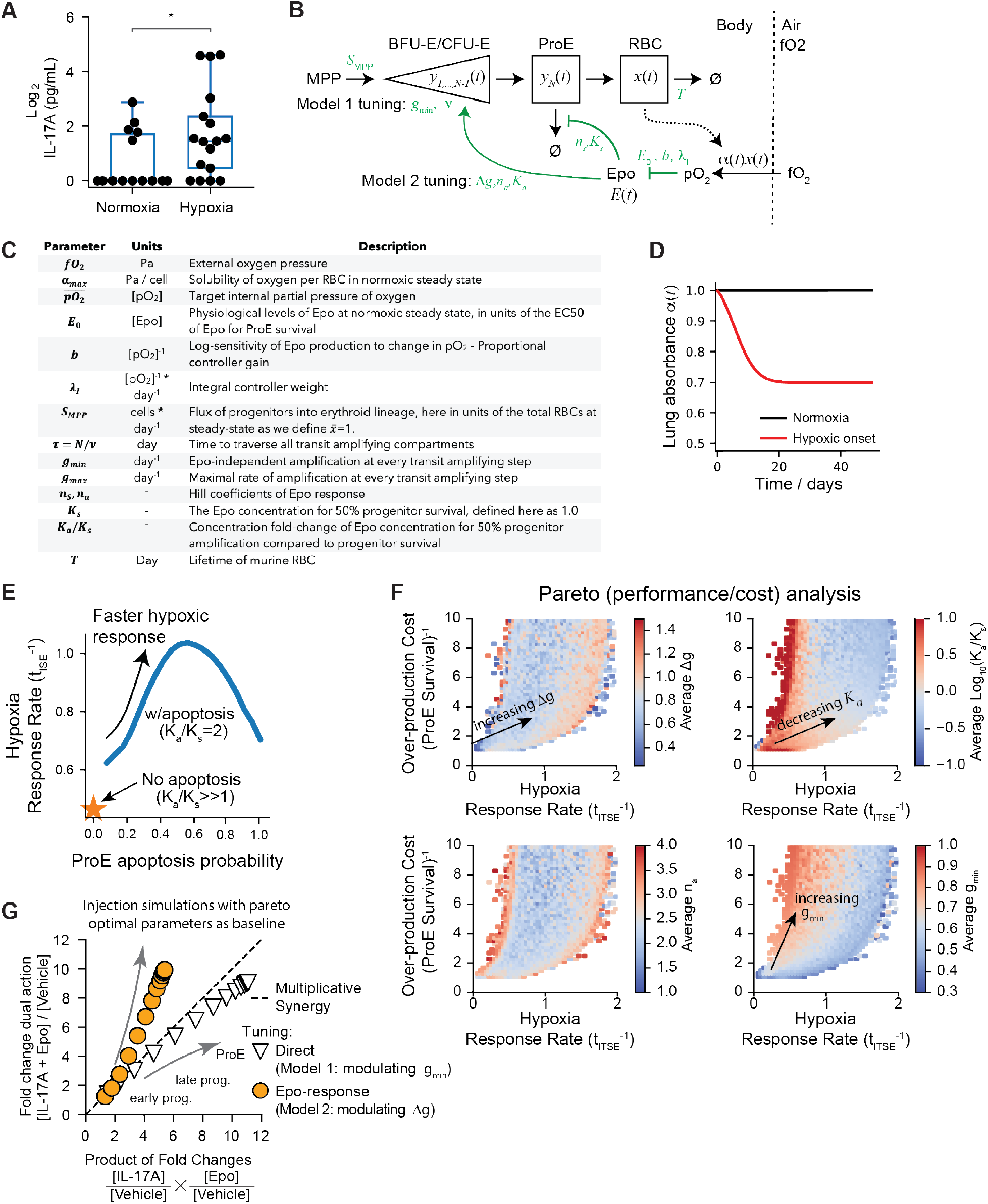
Modeling the cost-speed trade-off in erythropoietic feedback control. **A** Measured blood serum concentrations of IL-17A in mice in normoxia and following 24 hours of exposure to hypoxia following the treatment schema in **Fig. 3D**. Measurements are carried out by immunoPCR along with recombinant murine IL-17A as a standard curve. Data points correspond to individual mice and box plots are defined as in **Fig. 2B**. The increase in IL-17A in hypoxia is statistically significant (Wilcoxon rank-sum test, one-tailed p-value <0.05) but still well below the EC^50^ of erythroid progenitors seen in CFU-E assays. This measurement argues against IL-17A acting as a second hypoxia signal. **B** Diagram of the dynamical systems model of erythropoietic feedback control. Black arrows indicate cell flux. Green arrows indicate regulatory links. Symbols in green show parameters associated with different model components. The full mathematical model is defined in **Supplementary Text 1. C** Table defining the model parameters in (B). **D** The dynamics of lung absorbance used to model in disease-associated hypoxic onset in **Fig. 4B** and all subsequent analyses (**Fig. 4C-E**, and panels **E,F** below). The graphs show the value of model parameter for oxygen absorbance through the lung, *α*(*t*), as defined in **Supplementary Text 1** (Eq. [10]). **E** The model recapitulates a requirement for ProE apoptosis in accelerating hypoxic response rates, defined as in **Fig. 4B**. The orange star denotes the response rate when *K*_*a*_/*K*_*s*_ ≫ 1, *g*_*min*_ = 0.3/day, Δ*g* = 0.9/day, corresponding to a case where all ProE survive and Epo exclusively regulates cell proliferation. The blue curve corresponds to the same Epo regulation of progenitors, but now with *K*_*a*_/*K*_*s*_ = 2.0 corresponding to Epo regulating both response rate and survival. Variation along the blue curve represents varying values of the Epo-independent proliferation rate *g*_*min*_, with higher values leading to high apoptosis rates. **F** Random sampling of model parameters in 10^5^ simulations reveal different trade-offs in overproduction cost and response rate across parameters. This panel extends **Fig. 4E**. Parameters were sampled over the intervals: *g*_*min*_ ∈ [0.3,1.3]/day, Δ*g* ∈ [0.3,1.3]/day, K_*a*_/K_*s*_ ∈ [0.05,20], *n*_*a*_ ∈ [1,3]. **H** Computational predictions for the expansion of erythroid progenitors in each of the two Models following Epo + IL-17A treatment as compared to multiplicative fold change of Epo and IL-17A alone. Pareto optimal parameters were used to define baseline. The action of model 2 is defined here as modulation of Δ*g*. The dashed line indicates multiplicative synergy. Refer to **Supplementary Text 1** for modeling parameters used.

**Supplementary Figure 7.**
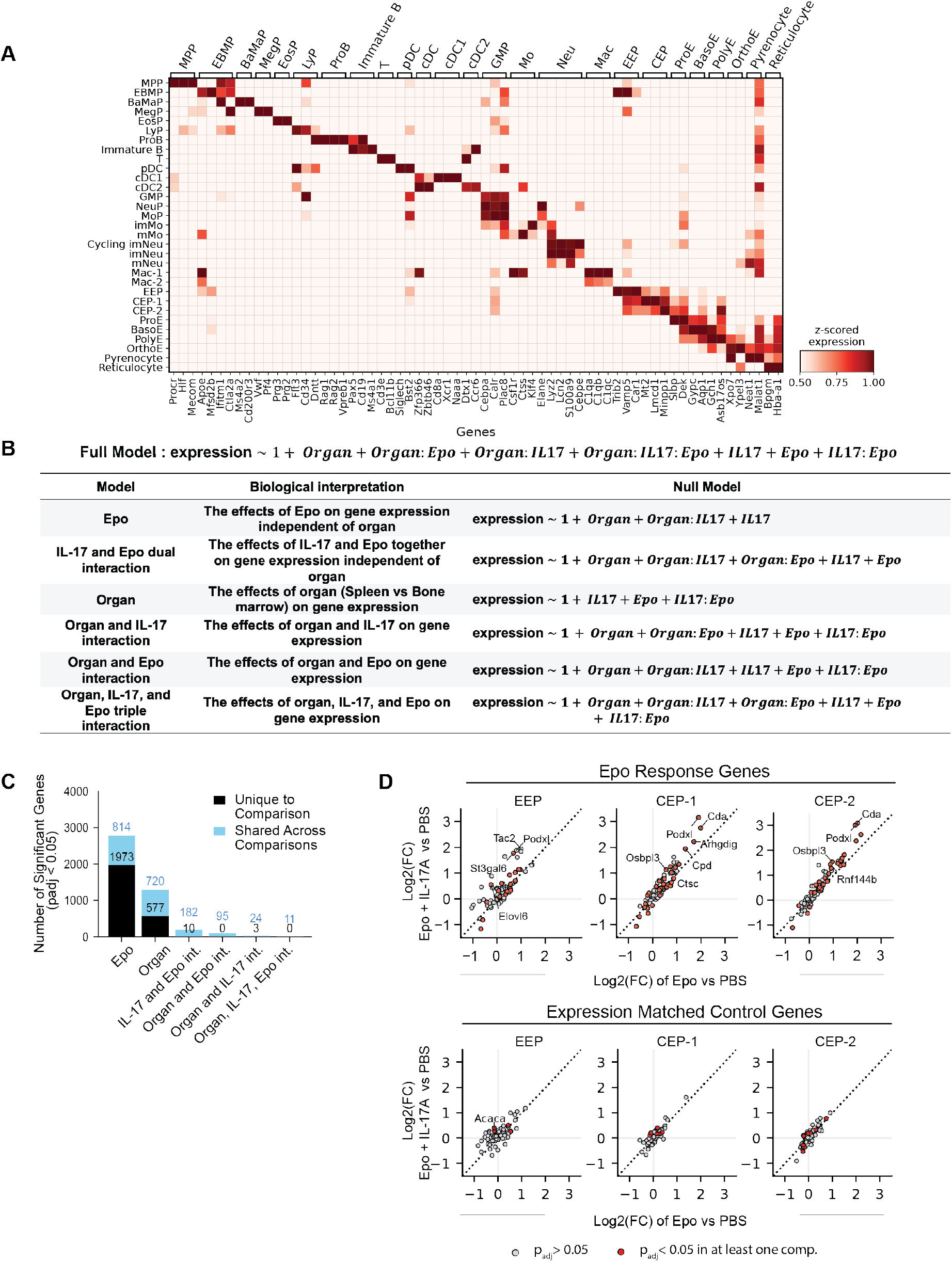
Annotation of cell states in the scRNA-Seq data and analysis of their transcriptional response. **A** Heatmap showing expression of cell type-specific marker genes (columns) across cell clusters annotated to different hematopoietic cell states (rows). The genes were selected from literature as markers of cell states, as shown by their group label at the top. Marker gene list and the corresponding origins can be found in **Table S4**. The diagonal pattern demonstrates marker gene specificity for their respective cell populations. Expression values are the mean of z-score standardized values. **B** Definition of the generalized linear model (GLM) and the nested null models used to dissect gene expression dependency on cytokine treatments and organ-of-origin for each gene, supporting **Fig. 5F**. The full model incorporates all possible interactions between organ (spleen vs bone marrow), Epo, and IL-17A treatments. The nested models remove terms to enable statistical significance testing for different terms (Likelihood Ratio Tests followed by multiple hypothesis correction across all genes, see Methods). The biological interpretation and corresponding null model formula are shown for each comparison. **C** Distribution of significantly regulated genes (p_adj_ < 0.05, |log_2_FC| > 0.25) from applying the GLM to cells annotated as CEP-1 (equivalent to early CFU-E). The six bars correspond to testing against each of the null models tabulated in (B). Black bars indicate genes uniquely regulated in a single comparison, while blue bars show genes shared across multiple comparisons. The analysis reveals that Epo treatment and tissue-specific differences are the dominant sources of transcriptional variation. The IL-17A and Epo interaction showed limited unique gene regulation (10 unique genes), with most regulated genes shared with other comparisons. Similar distributions were observed across other erythroid populations. The number of genes which are overlapping between Epo treatment and IL-17A and Epo interaction are shown in **Fig. 5F. D** Top: comparison of changes in the expression of Epo response genes across erythroid progenitor states. Red dots indicate genes with significant differential expression (p_adj_ < 0.05) in either Epo+IL-17A vs PBS or Epo vs PBS comparisons. These Epo response genes (**Table S4**, from Tusi et al.^13^) were used to calculate response scores in **Fig. 6A**. Select genes are labeled. Dotted lines indicate y=x. Log2(FC) = log2(fold-change). Bottom: the same analysis repeated for randomly-selected expression-matched control genes shows that the increase seen in Epo response is specific.

## Supplemental text: Dynamical modeling of a burden/response trade-off in erythropoiesis

This supplement defines the dynamical model used to evaluate alternative hypotheses for homeostatic control tuning of red blood cell (RBC) production, supporting **Figs. 4** and **S6**.

### 1. Model Definition

We adopt a dynamical systems model of erythropoiesis represented by a series of ODEs and algebraic equations describing system variables. Several prior ODE models of erythropoiesis have been developed^1^ that incorporate a considerably greater level of detail than described here, but these lacked the control elements of the current model.

#### 1.1 Model design overview

The model is designed to capture the relevant biological processes that effect a trade-off between two performance goals of the erythropoietic homeostatic control circuit, and which are likely under control of an early-expressed receptor: (i) maximizing the speed of recovery from hypoxic stress, and (ii) minimizing the burden of constitutive progenitor cell over-production in normoxia. This trade-off results from two key features of RBC production:

1. A lag time in differentiation and the transit amplification of erythroid progenitors^2,3^.
2. A homeostatic negative feedback circuit: red blood cells (RBCs) elevate oxygen blood concentration (pO_2_); pO_2_ represses Epo production; and Epo stimulates RBC production ^4,5^.

We incorporate two specific the mechanisms of Epo action on erythropoiesis: (1) Epo prevents death of later precursors (proerythroblasts, or ProE) ^6–8^ ; and (2) Epo increases proliferation in progenitors^9,10^ leading to increased reticulocyte production in 5-7 days^11,12^, but does so at lower affinity.

The model does not incorporate many more detailed features of erythropoiesis, which we reasoned would add to complexity without qualitatively altering the key predictions that differentiate hypotheses regarding IL-17A function evaluated in the text. Notably, we ignore: (1) proliferation subsequent to ProE; (2) distinction between reticulocytes and RBCs; (3) nomenclature of early progenitors; (4) Epo uptake by erythroid progenitors; (5) density-dependent Epo responses mediated by Fas/FasL^10^. In addition, (6) we have not explicitly accounted for other effects of elevated Epo concentrations on erythropoiesis. Epo stimulates the release of premature reticulocytes into blood, offering a short-term response to hypoxic stress in <1 day ^13,14^. It additionally promotes division of late erythroid progenitors (EryA/B). These effects will quantitatively accelerate hypoxic responses, and all reducing resting costs and buffer the lag in generating new cells through progenitor division. However, they should do not qualitatively change RBC response profiles over longer periods (>3 days in mice, or >5-7 days in humans). The discriminating features related to IL-17A action discussed in this paper relate to longer latency periods and we reasoned that they can be described without incorporating the late-stage behaviors.

#### 1.2 Model variables

The model, shown schematically in **Figs. 4A, S6B**, has the following dynamic variables:

| Variable | Units | Description |
| --- | --- | --- |
| $pO_2(t)$ | Pa | Partial pressure of oxygen (in blood) |
| $E(t)$ | - | Erythropoietin concentration, in units of the EC50 for ProE survival |
| $y_{1,...,N-1}(t)$ | # cells | Number of progenitor cells along $N-1$ steps of differentiation |
| $y_N(t)$ | # cells | Number of ProE cells |
| $x(t)$ | # cells | Number RBCs |

These variables include *N* numerically labeled stages of progenitor differentiation, where a choice of *N* is used to limit the variance in the latency time of transit amplification. The choice of *N* is not intended to reflect specific biological transitions, although one may think of BFU-E/EEP as reflecting early progenitors *y*_1,…,*n*<*N*−1_, and CFU-E/CEP as reflecting the later pro-genitors *y*_*n*+1,…,*N*−1_. Variation in *N* is not expected to alter model conclusions, provided *N* ≫ 1. We have used *N*=20. A more precise definition of “large N” is given in section 3 below. For the current analysis, the transition stage *n* between EEP and CEP plays has no role and is not discussed further.

#### 1.3 Model equations and parameters

The model architecture is shown schematically in **Fig. S6B** and is fully defined in the following sub-sections. The model parameters are non-dimensionalized where possible as described in the tables below.

##### 1.3.1 Fast variables: oxygen and erythropoietin

We assume that oxygen and Epo concentrations (*pO*_2_(*t*), *E*(*t*)) are in quasi-steady-state as they equilibrate rapidly compared to the timescale of cell differentiation in erythropoiesis. Oxygen equilibration occurs within minutes through pulmonary gas exchange. Epo half-life in the blood is ∼12 hours. By contrast, the dynamics of cell populations responds over days.

In our model the oxygen dissolved in blood is assumed to be directly proportional to the RBC mass, i.e.:

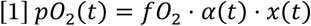

where here *fO*_2_ is the environmental oxygen pressure, and *α*(*t*) is the solubility of oxygen per RBC at time *t*. In steady-state healthy conditions, we assume *α*(*t*) = *α*_*max*_. Simulations of infection-induced hypoxic stress are carried out by providing a dynamic value of *α*(*t*) that drops from *α*_*max*_ over time to a lower value, as discussed below. A more detailed model of oxygen transport would address changes in the cooperativity of hemoglobin as a function of blood pH, as well as changes in hematocrit. Both vary with hypoxia. However, in this model we make a simplification by ignoring cooperativity of binding and treated RBC mass rather than blood concentration.

For Epo, production is generated in response to hypoxia by cells in the kidney and we take it to be:

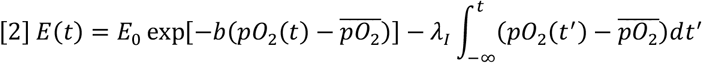

Here, the first term reflects proportional control of Epo production in the kidney in response to changes in oxygen tension. Eq. [2] is defined such that *E*(*t*) = *E*_0_ at steady-state. The concentration of Epo mRNA has been shown to grow exponentially with the drop in partial pressure, which is reflected in the exponential factor with sensitivity to oxygen changes *b*. In addition, we include a second term reflecting an integral feedback controller with strength *λ*_*I*_ [Eqn. 2]. Integral feedback control ensures robustness of the steady-state to fluctuations in system parameters. In summary, the table below summarizes the parameters included in this sub-section:

| Parameter | Units | Description |
| --- | --- | --- |
| $fO_2$ | Pa | External oxygen pressure |
| $\alpha_{max}$ | Pa / cell | Solubility of oxygen per RBC in normoxic steady state |
| $\overline{pO_2}$ | [pO<sub>2</sub>] | Target internal partial pressure of oxygen |
| $E_0$ | [Epo] | Physiological levels of Epo at normoxic steady state, in units of the EC50 of Epo for ProE survival |
| $b$ | [pO<sub>2</sub>]<sup>-1</sup> | Log-sensitivity of Epo production to change in pO<sub>2</sub> - Proportional controller gain |
| $\lambda_I$ | [pO<sub>2</sub>]<sup>-1</sup> * day<sup>-1</sup> | Integral controller weight |

##### 1.3.2 Slow variables: erythroid progenitors

**Figs. 4A, S6B** show schematically the differentiation hierarchy from multipotent progenitors to RBCs. This hierarchy is model through a set of coupled linear ODEs:

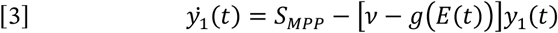

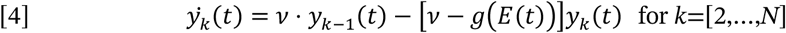

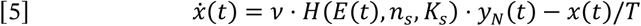

Here, Eqs. [3] and [4] describe the dynamics of the BFU-E and CFU-E cells progressing through stages of differentiation, and Eq. [5] describes dynamics of the RBCs.

Eq. [3] describes dynamics of the earliest erythroid-committed progenitors [compartment *y*_1_], which arise from progenitors (MPPs) that are not under control of Epo. The rate of MPP differentiation is thus treated as constant in the model, defined by the rate parameter *S*_*MPP*_. In both Eqs. [3] and [4], progenitors in each compartment *y*_*k*_ differentiate to the subsequent compartment *y*_*k*+1_ with mean rate *ν*. The progenitors proliferate with an Epo-dependent rate *g*(*E*), given by

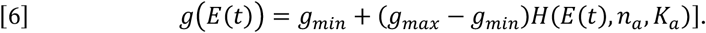

where *g*_*min*_ is the Epo-independent early progenitor proliferation rate, *g*_*max*_ is the maximal proliferation rate at saturating Epo concentrations, and *H*(*z, n, K*) = *z*^*n*^/[*K*^*n*^ + *z*^*n*^] is a Hill function with coefficient *n* and half-max concentration *K*.

The dynamics of RBCs (Eq. [5]) incorporates the final elements of the model: the first term in this equation describes ProE cells [compartment *y*_*N*_] differentiating into RBCs at a rate *ν*, but with only a fraction *H*(*E*(*t*), *n*_*s*_, *K*_*s*_) of the cells surviving in an Epo-dependent manner. Here *H* is a Hill function defined as above, but with a different half-max concentration *K*_*s*_ for ProE survival and coefficient *n*_*s*_. RBCs are cleared with an average lifetime *T*. As discussed in subsection 1.1, this model ignores cell division after ProE formation and does not distinguish reticulocytes from RBCs.

The model parameters in Eqs. [3-6] are summarized in the following table, along with the values used for simulation.

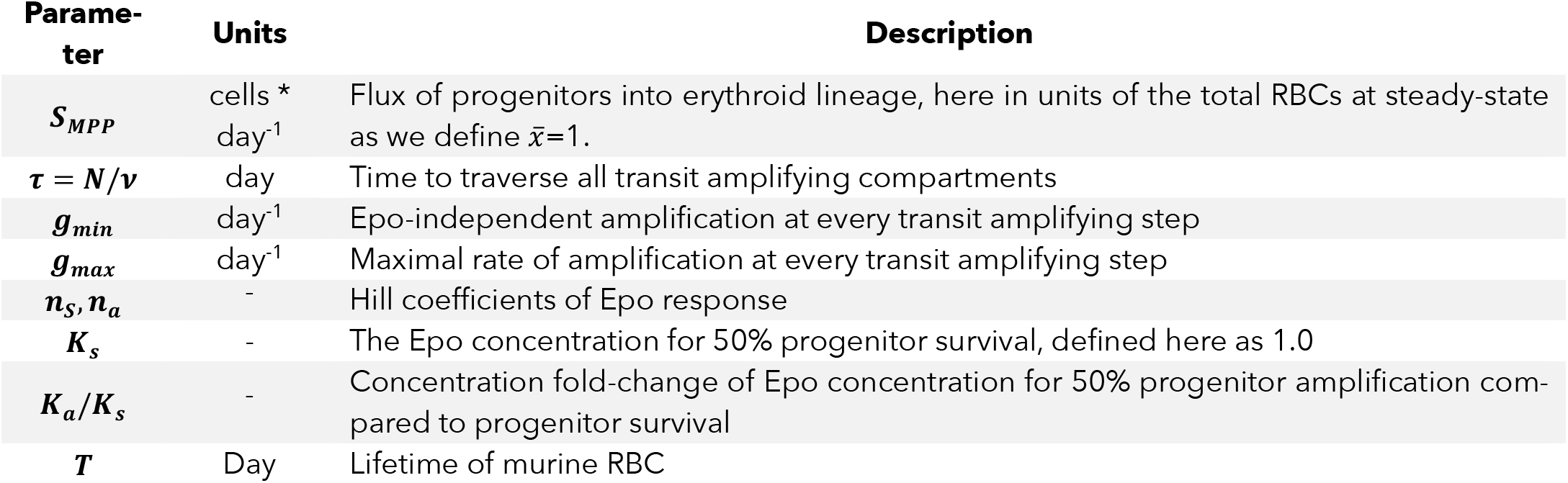

### 2. Normoxic steady state and its constitutive over-production cost

The over-production of ProE in steady state normoxic conditions represents a constitutive cost of the model. We define the over-production burden as the ratio of produced ProE cells to those surviving, i.e., cost *θ*_*c*_

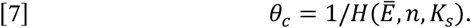

To calculate this cost for a given parameter choice, we solve the dynamical equations at steady state with *α*(*t*) = *α*_*max*_ = 1. We denote steady-state values of the dynamical system variables with an over-bar (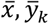, etc). At steady-state, Eq. [2] enforces 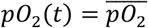, and so from Eq. [1] the steady-state RBC mass is, 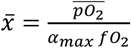. A convenient choice of units sets, 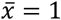 as discussed in section 3 below.

We now provide the steady-state solutions for the cell populations, 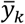 from Eqs. [3-6]:

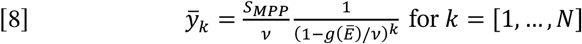

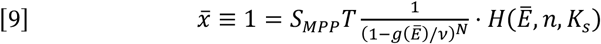

A steady-state is achieved when the parameters thus satisfy Eq. [9]. When *N* is large,, 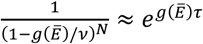 and is independent of *N*. Note that *Ē* only enters the steady-state equations through its EC50 values *Ē*/*K*_*s*_, *Ē*/*K*_*a*_. Its value is not determined by Eq. [2] because the integral control term allows adding an arbitrary offset *Ē* = *E*_0_ + *δE*, e.g., 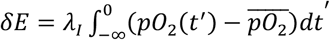.

In this study, a numerical solver was used in Python to solve Eq. [8] for Ē= once remaining model parameters were established. For simplicity, we set *E*_0_ = Ē = for a given choice of parameters. Subsequently, the over-production cost *θ*_*c*_ can be calculated from Eq. [9].

### 3. Parameter values

Parameter values used in simulation are defined in the tables above, with justification provided here. Without loss of generality, we have set units of oxygen pressure and cell number such that:

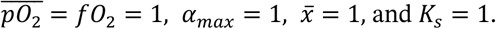

And *E*_0_ is set by solving the steady-state equation Eq. [9] as described above. For the remaining parameters, we established values as follows.

1. The lifetime of RBCs in mice has been estimated several times in the literature with broadly consistent results. We have used the value of *T=*40 days (see reference in table above).
2. The time for mouse BFU-Es to give rise to colonies containing RBC is ∼7 days, and so we have set *τ* = *N*/*ν* = 7 days.
3. The choice of *N* is constrained by two technical aspects of the ODE model. We have used a choice of *N* = 20. We motivate our choice as follows.
  a. In Eq. [9], as noted the solution is independent of *N* when *N* is large, so the precise value is not important.
  b. In an ODE modeling framework, the time taken to transition through a chain of *N* steps with equal transition rates *ν* follows a gamma distribution *t*∼Gamma,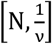. As such *N* sets the variability in the time of transit amplification. One notes that for the gamma distribution the mean time is 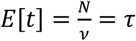, and standard deviation is,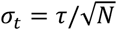. Therefore, *N* may be thought of as a convenience variable used to tune the sharpness of the latency time of the model, with a coefficient of variation,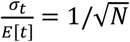.
4. From Eq. [9] and assumptions above, we note that, 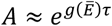 corresponds to the total amplification that progenitors undergo. The precise number of cell divisions that progenitors undergo is not measured but is thought to be ∼10. We have chosen an amplification of *A* = 1000. From this we can constrain *g*(Ē =)∼0.8/day, which sets upper and lower bounds on *g*_*min*_ and *g*_*max*_. This estimate is also in agreement with prior measurements of the mean cell division time for BFU-E/CFU-E in the fetal liver of mice, which is ∼15 hours. The doubling time is, 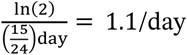1.1/day.
5. Only one parameter remains that governs the steady-state behavior: the rate of cells entering the erythroid lineage from the MPP pool *S*_*MPP*_. From Eq. [9] and assumptions 1-3,5 above, we can relate these parameters *S*_*MPP*_ = (1 − *g*_*min*_*τ*/*N*)^−*N*^*θ*_*c*_/*T*. We note that (1 − *g*_*min*_*τ*/*N*)^−*N*^ = *A* corresponds to the total amplification that progenitors undergo as defined in assumption 4 above. Thus,, 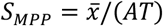 is fully specified.
6. We assume that, in the absence of IL-17A, the early progenitors are less responsive to Epo and thus *K*_*a*_ > *K*_*s*_. (Note that *K*_*s*_ = 1 sets the units of Epo, as discussed above).

This now leaves three parameters unconstrained but bounded: *K*_*a*_/*K*_*s*_, *g*_*min*_, *g*_*max*_. The results given in the paper hold for different choices of these three parameters. As a baseline, the following values were used:

### 4. Model Simulations.

#### 4.1 Simulations (Figs. 4B-D, S6C)

The dynamical equations were implemented and solved using Python v3.10.13, with the code available at github.com/AllonKleinLab/Wu2024. We used the `integrate.solve_ivp` function from SciPy v1.11.4, applying the ‘BDF’ method to solve the equations. Unless stated otherwise, model parameters used are as listed in Table I.

**Table I.** Baseline parameter values.

| Parameter | Value | Estimate Range | Reference |
| --- | --- | --- | --- |
| $\overline{pO_2}$ | 1.0 | Units set to 1.0 | - |
| $fO_2$ | 1.0 | Units set to 1.0 | - |
| $E_0$ | Set by Eq. [9] | - | - |
| $\alpha_{max}$ | 1.0 | - | - |
| $S_{MPP}$ | $1.25 \cdot 10^{-4} / \text{day}$ | - | - |
| $\tau$ | 7.0 days | 7 Days | 12,15 |
| $g_{min}$ | 0.5/day | - | - |
| $g_{max}$ | 0.8/day | 1-3/day | - |
| $n_a, n_s$ | 2.0 | 1-3 | 16 |
| $K_s$ | 1.0 | Units set to 1.0 | - |
| $K_a/K_s$ | 1.5 | Lower Epo sensitivity in progenitors than ProE | 17,18 |
| $T$ | 40.0 days | $40 \pm 1.9$ days | 19 |
| $b$ | 5 | - | - |
| $\lambda_I$ | 0.05 | - | - |

For the dynamics, we numerically solved the system dynamics with a model of deteriorating pulmonary function, modeled as a biphasic decay in Eq. [1] with:

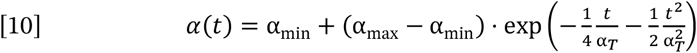

Here *α*(0) = α_max_ = 1.0 is the absorbance in the healthy initial state, *α*(*t* ≫ α_*T*_) = α_min_ is the terminal absorbance, and α_*T*_ is the timescale for lung function to drop to approximately the midpoint. In all simulations we used α_*T*_ = 5 days and α_*min*_ = 0.7, indicating that the lung’s absorbance is only 70% of healthy lung capacity.

#### 4.2 Defiinition ofi perfiormance goals

From the numerical solution of the model response to deteriorating pulmonary function (Eq. [10]), we define a time-scale for recovery by adapting the integral-time-square error (ITSE), a common metric for control system performance:

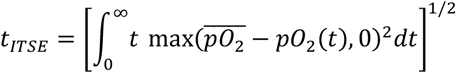

For every parameter set considered, one can now define two performance goals: the response performance *s*=1/*t*_*ITSE*_, and the constitutive over-production burden *θ*_*c*_ (Eq. [7]).

#### 4.3 Parameter scanning and response/burden (s, θ_c_) trade ofifi analysis

Here we describe the parameter scans shown in **Figs. S6F, 4D, E**. These figures plot the performance goals (*s, θ*_*c*_) for different parameter sets.

##### 4.3.1 Single-parameter scans (Figs. 4D, S6E)

In **Fig. 4D**, performance goals are evaluated while varying one parameter at a time of the ranges shown in **Table II**.

**Table II:** Parameter ranges scanned.

| Table II: Parameter ranges scanned. |  |
| --- | --- |
| Parameter | Range |
| $\Delta g$ | 0.3-1.3/day |
| $K_a/K_s$ | 0.05 - 20 |
| $n_a$ | 1 - 4 |
| $g_{\min}$ | 0.3 - 1.3/day |
| $\tau$ | 5-11 days |
| $S_{MPP}$ | $(6.25 - 62.5) \cdot 10^{-5}/\text{day}$ |

**Table III:** Parameter ranges scanned.

| Parameter | Range |
| --- | --- |
| $\Delta g$ | 0.25-1.5/day |
| $K_a/K_s$ | 0.05 - 20 |
| $n_a$ | 1 - 4 |
| $g_{min}$ | 0.3 – 1.0/day |

Using baseline parameters, the scan for *g*_*min*_ was then used to construct **Fig. S6E**, showing the relationship between performance *s* and apoptosis rate (1 − 1/*θ*_*c*_).

##### 4.3.2 Multi-parameter sampling (Fig. 4E, S6F)

For **Fig. 4E, S6F**, we sampled 10^5^ parameters sets (*g*_*min*_, Δ*g, K*_*a*_/*K*_*s*_, *n*_*a*_) on a latin hypercube. Latin hypercube sampling was performed using subroutine *lhs* from python package pyDOE2 (v1.3.0). For each parameter value we then calculated the performance goals (*s, θ*_*c*_). The simulations were binned over (*s, θ*_*c*_) values, and the mean value of each parameter in each bin was plotted. The values of *g*_*min*_, Δ*g*, *K*_*a*_/*K*_*s*_ given in the baseline table above are close to the pareto front.

All remaining parameters have values as in parameters have values as in **Table I**.

##### 4.4 Simulation ofi experimental injections under competing models ofi IL-17A action (Fig. 4F, S6G)

The synergy plots shown in **Fig. 4F** and **Fig. S6G** provide a qualitative distinction between the two models. The simplifications made in model construction (see introduction and section 1 above) make this model unsuitable for *bona fiide* data fitting and just one choice of parameters is sufficient to highlight qualitative differences between the hypotheses.

To generate the synergy plots shown in **Fig. 4F** and **Fig. S6G** we numerically solved the system dynamics (Eqs. [1-5]) as described in section 4.1 over the time interval *t=*[0,3] days and using the same baseline parameters (Table I), with the following modifications:

1. We held *α*(*t*) = *α*_*max*_ = constant.
2. To simulate Epo injection, we replaced Eq. [2] with the following:

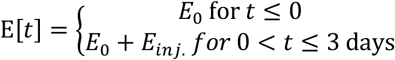

with *E*_*inj*._ = 5*K*_*s*_

To simulate IL-17A injection, we acutely changed the model parameters as follows:

1. For Model 1 (direct control):

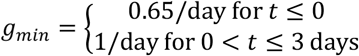
2. For Model 2 (Epo-sensitization)

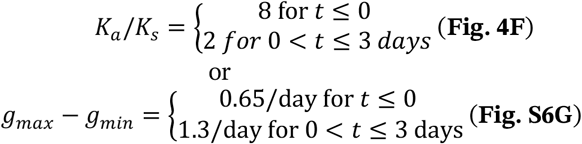

For joint injections, both (2) and (3) were implemented. In each case, the dynamics were solved, and data points plotted correspond to values ([*y*_k_]^(E)^[*y*_k_]^(I)^/([K*y*]^(0)^)^2^, [*y*_k_]^(E+I)^/[*y*_k_]^(0)^) for increasing values of *k* at time *t*=3 days post-injection.

## References

1. Ziegler, J.G., and Nichols, N.B. (2022). Optimum Settings for Automatic Controllers. Transactions of the American Society of Mechanical Engineers 64, 759–765. 10.1115/1.4019264.

2. Åström, K.J., and Hägglund, T. (2006). Advanced PID control (International Society of Automation).

3. Kitano, H. (2004). Biological robustness. Nat Rev Genet 5, 826–837.

4. Szekely, P., Sheftel, H., Mayo, A., and Alon, U. (2013). Evolutionary tradeoffs between economy and effectiveness in biological homeostasis systems. PLoS Comput Biol 9, e1003163. 10.1371/journal.pcbi.1003163.

5. Shoval, O., Sheftel, H., Shinar, G., Hart, Y., Ramote, O., Mayo, A., Dekel, E., Kavanagh, K., and Alon, U. (2012). Evolutionary trade-offs, Pareto optimality, and the geometry of phenotype space. Science (New York, N.Y.) 336, 1157–1160. 10.1126/science.1217405.

6. Romanovsky, A.A. (2007). Thermoregulation: some concepts have changed. Functional architecture of the thermoregulatory system. Am J Physiol Regul Integr Comp Physiol 292, R37–46. 10.1152/ajpregu.00668.2006.

7. Saltiel, A.R., and Kahn, C.R. (2001). Insulin signalling and the regulation of glucose and lipid metabolism. Nature 414, 799–806. 10.1038/414799a.

8. Brown, E.M. (1991). Extracellular Ca2+ sensing, regulation of parathyroid cell function, and role of Ca2+ and other ions as extracellular (first) messengers. Physiol Rev 71, 371–411. 10.1152/physrev.1991.71.2.371.

9. Sender, R., Fuchs, S., and Milo, R. (2016). Revised Estimates for the Number of Human and Bacteria Cells in the Body. PLoS Biol 14, e1002533. 10.1371/journal.pbio.1002533.

10. Koulnis, M., Porpiglia, E., Hidalgo, D., and Socolovsky, M. (2014). Erythropoiesis: From Molecular Pathways to System Properties. In A Systems Biology Approach to Blood, S.J. Corey, M. Kimmel, and J.N. Leonard, eds. (Springer New York), pp. 37–58. 10.1007/978-1-4939-2095-2_3.

11. Erslev, A. (1953). Humoral regulation of red cell production. Blood 8, 349–357.

12. Erslev, A.J. (1997). Clinical erythrokinetics: a critical review. Blood Rev 11, 160-167. S0268-960X(97)90011-4 [pii].

13. Tusi, B.K., Wolock, S.L., Weinreb, C., Hwang, Y., Hidalgo, D., Zilionis, R., Waisman, A., Huh, J.R., Klein, A.M., and Socolovsky, M. (2018). Population snapshots predict early haematopoietic and erythroid hierarchies. Nature 555, 54–60. 10.1038/nature25741.

14. Palis, J. (2014). Primitive and definitive erythropoiesis in mammals. Frontiers in Physiology 5. 10.3389/fphys.2014.00003.

15. Pop, R., Shearstone, J.R., Shen, Q., Liu, Y., Hallstrom, K., Koulnis, M., Gribnau, J., and Socolovsky, M. (2010). A key commitment step in erythropoiesis is synchronized with the cell cycle clock through mutual inhibition between PU.1 and S-phase progression. PLoS Biol 8.

16. Jacobson, L.O., Goldwasser, E., Fried, W., and Plzak, L. (1957). Role of the kidney in erythropoiesis. Nature 179, 633–634. 10.1038/179633a0.

17. Takeda, K., Aguila, H.L., Parikh, N.S., Li, X., Lamothe, K., Duan, L.J., Takeda, H., Lee, F.S., and Fong, G.H. (2008). Regulation of adult erythropoiesis by prolyl hydroxylase domain proteins. Blood 111, 3229–3235. 10.1182/blood-2007-09-114561.

18. Koury, M.J., and Bondurant, M.C. (1990). Erythropoietin retards DNA breakdown and prevents programmed death in erythroid progenitor cells. Science (New York, N.Y.) 248, 378–381.

19. Koulnis, M., Porpiglia, E., Porpiglia, P.A., Liu, Y., Hallstrom, K., Hidalgo, D., and Socolovsky, M. (2012). Contrasting dynamic responses in vivo of the Bcl-xL and Bim erythropoietic survival pathways. Blood 119, 1228-1239. blood-2011-07-365346 [pii] 10.1182/blood-2011-07-365346.

20. Koulnis, M., Liu, Y., Hallstrom, K., and Socolovsky, M. (2011). Negative autoregulation by Fas stabilizes adult erythropoiesis and accelerates its stress response. PLoS One 6, e21192. 10.1371/journal.pone.0021192PONE-D-11-05396[pii].

21. Hidalgo, D., Bejder, J., Pop, R., Gellatly, K., Hwang, Y., Maxwell Scalf, S., Eastman, A.E., Chen, J.J., Zhu, L.J., Heuberger, J., et al. (2021). EpoR stimulates rapid cycling and larger red cells during mouse and human erythropoiesis. Nature communications 12, 7334. 10.1038/s41467-021-27562-4.

22. Wu, H., Liu, X., Jaenisch, R., and Lodish, H.F. (1995). Generation of committed erythroid BFU-E and CFU-E progenitors does not require erythropoietin or the erythropoietin receptor. Cell 83, 59–67.

23. Gregory, C.J. (1976). Erythropoietin sensitivity as a differentiation marker in the hemopoietic system: studies of three erythropoietic colony responses in culture. Journal of cellular physiology 89, 289–301. 10.1002/jcp.1040890212.

24. Peschle, C., Migliaccio, A.R., Migliaccio, G., Ciccariello, R., Lettieri, F., Quattrin, S., Russo, G., and Mastroberardino, G. (1981). Identification and characterization of three classes of erythroid progenitors in human fetal liver. Blood 58, 565–572.

25. Porpiglia, E., Hidalgo, D., Koulnis, M., Tzafriri, A.R., and Socolovsky, M. (2012). Stat5 signaling specifies basal versus stress erythropoietic responses through distinct binary and graded dynamic modalities. PLoS Biol 10, e1001383. 10.1371/journal.pbio.1001383PBIOLOGY-D-12-00220[pii].

26. Bauer, A., Tronche, F., Wessely, O., Kellendonk, C., Reichardt, H.M., Steinlein, P., Schutz, G., and Beug, H. (1999). The glucocorticoid receptor is required for stress erythropoiesis. Genes Dev 13, 2996–3002.

27. Broudy, V.C. (1997). Stem cell factor and hematopoiesis. Blood 90, 1345–1364.

28. Lenox, L.E., Perry, J.M., and Paulson, R.F. (2005). BMP4 and Madh5 regulate the erythroid response to acute anemia. Blood 105, 2741–2748.

29. Perry, J.M., Harandi, O.F., and Paulson, R.F. (2007). BMP4, SCF, and hypoxia cooperatively regulate the expansion of murine stress erythroid progenitors. Blood 109, 4494–4502. blood-2006-04-016154 [pii] 10.1182/blood-2006-04-016154.

30. Cua, D.J., and Tato, C.M. (2010). Innate IL-17-producing cells: the sentinels of the immune system. Nat Rev Immunol 10, 479–489. 10.1038/nri2800.

31. Gaffen, S.L. (2009). Structure and signalling in the IL-17 receptor family. Nat Rev Immunol 9. 10.1038/nri2586.

32. Li, X., Bechara, R., Zhao, J., McGeachy, M.J., and Gaffen, S.L. (2019). IL-17 receptor-based signaling and implications for disease. Nat Immunol 20, 1594–1602. 10.1038/s41590-019-0514-y.

33. Veldhoen, M. (2017). Interleukin 17 is a chief orchestrator of immunity. Nat Immunol 18, 612–621. 10.1038/ni.3742.

34. Wright, J.F., Bennett, F., Li, B., Brooks, J., Luxenberg, D.P., Whitters, M.J., Tomkinson, K.N., Fitz, L.J., Wolfman, N.M., Collins, M., et al. (2008). The human IL-17F/IL-17A heterodimeric cytokine signals through the IL-17RA/IL-17RC receptor complex. J Immunol 181, 2799–2805.

35. Socolovsky, M., Nam, H., Fleming, M.D., Haase, V.H., Brugnara, C., and Lodish, H.F. (2001). Ineffective erythropoiesis in Stat5a(-/-)5b(-/-) mice due to decreased survival of early erythroblasts. Blood 98, 3261–3273.

36. Liu, Y., Pop, R., Sadegh, C., Brugnara, C., Haase, V.H., and Socolovsky, M. (2006). Suppression of Fas-FasL coexpression by erythropoietin mediates erythroblast expansion during the erythropoietic stress response in vivo. Blood 108, 123–133.

37. Swaminathan, A., Hwang, Y., Winward, A., and Socolovsky, M. (2022). Identification and Isolation of Burst-Forming Unit and Colony-Forming Unit Erythroid Progenitors from Mouse Tissue by Flow Cytometry. J Vis Exp. 10.3791/64373.

38. El Malki, K., Karbach, S.H., Huppert, J., Zayoud, M., Reißig, S., Schüler, R., Nikolaev, A., Karram, K., Münzel, T., Kuhlmann, C.R.W., et al. (2013). An Alternative Pathway of Imiquimod-Induced Psoriasis-Like Skin Inflammation in the Absence of Interleukin-17 Receptor A Signaling. Journal of Investigative Dermatology 133, 441–451. 10.1038/jid.2012.318.

39. Conti, H.R., and Gaffen, S.L. (2015). IL-17–Mediated Immunity to the Opportunistic Fungal Pathogen Candida albicans. The Journal of Immunology 195, 780–788. 10.4049/jimmunol.1500909.

40. Iwakura, Y., Ishigame, H., Saijo, S., and Nakae, S. (2011). Functional specialization of interleukin-17 family members. Immunity 34, 149–162. 10.1016/j.immuni.2011.02.012.

41. Kumar, P., Monin, L., Castillo, P., Elsegeiny, W., Horne, W., Eddens, T., Vikram, A., Good, M., Schoenborn, A.A., Bibby, K., et al. (2016). Intestinal Interleukin-17 Receptor Signaling Mediates Reciprocal Control of the Gut Microbiota and Autoimmune Inflammation. Immunity 44, 659–671. 10.1016/j.immuni.2016.02.007.

42. de Boer, J., Williams, A., Skavdis, G., Harker, N., Coles, M., Tolaini, M., Norton, T., Williams, K., Roderick, K., Potocnik, A.J., and Kioussis, D. (2003). Transgenic mice with hematopoietic and lymphoid specific expression of Cre. European Journal of Immunology 33, 314–325. 10.1002/immu.200310005.

43. Heinrich, A.C., Pelanda, R., and Klingmuller, U. (2004). A mouse model for visualization and conditional mutations in the erythroid lineage. Blood 104, 659–666.

44. Kato, M., Kato, Y., and Sugiyama, Y. (1999). Mechanism of the upregulation of erythropoietin-induced uptake clearance by the spleen. Am J Physiol 276, E887–895. 10.1152/ajpendo.1999.276.5.E887.

45. Hara, H., and Ogawa, M. (1976). Erthropoietic precursors in mice with phenylhydrazine-induced anemia. Am J Hematol 1, 453–458.

46. Goris, H., Bungart, B., Loeffler, M., Schmitz, S., and Nijhof, W. (1990). Migration of stem cells and progenitors between marrow and spleen following thiamphenicol treatment of mice. Exp Hematol 18, 400–407.

47. Colgan, S.P., Campbell, E.L., and Kominsky, D.J. (2016). Hypoxia and Mucosal Inflammation. Annual Review of Pathology: Mechanisms of Disease 11, 77–100. 10.1146/annurev-pathol-012615-044231.

48. Kumar, R., Singh, A.K., Starokadomskyy, P., Luo, W., Theiss, A.L., Burstein, E., and Venuprasad, K. (2021). Cutting Edge: Hypoxia-Induced Ubc9 Promoter Hypermethylation Regulates IL-17 Expression in Ulcerative Colitis. The Journal of Immunology 206, 936–940. 10.4049/jimmunol.2000015.

49. Konieczny, P., Xing, Y., Sidhu, I., Subudhi, I., Mansfield, K.P., Hsieh, B., Biancur, D.E., Larsen, S.B., Cammer, M., Li, D., et al. (2022). Interleukin-17 governs hypoxic adaptation of injured epithelium. Science (New York, N.Y.) 377, eabg9302. 10.1126/science.abg9302.

50. Dang, Eric V., Barbi, J., Yang, H.-Y., Jinasena, D., Yu, H., Zheng, Y., Bordman, Z., Fu, J., Kim, Y., Yen, H.-R., et al. (2011). Control of TH17/Treg Balance by Hypoxia-Inducible Factor 1. Cell 146, 772-784. 10.1016/j.cell.2011.07.033.

51. Boos, C.J., Woods, D.R., Varias, A., Biscocho, S., Heseltine, P., and Mellor, A.J. (2015). High Altitude and Acute Mountain Sickness and Changes in Circulating Endothelin-1, Interleukin-6, and Interleukin-17a. High Altitude Medicine & Biology 17, 25–31. 10.1089/ham.2015.0098.

52. Ritchie, N.D., Ritchie, R., Bayes, H.K., Mitchell, T.J., and Evans, T.J. (2018). IL-17 can be protective or deleterious in murine pneumococcal pneumonia. PLOS Pathogens 14, e1007099. 10.1371/journal.ppat.1007099.

53. Ndoricyimpaye, E.L., Van Snick, J., Robert, R., Bikorimana, E., Majyambere, O., Mukantwari, E., Nshimiyimana, T., Mbonigaba, V., Coutelier, J.P., and Rujeni, N. (2023). Cytokine Kinetics during Progression of COVID-19 in Rwanda Patients: Could IL-9/IFNγ Ratio Predict Disease Severity? Int J Mol Sci 24. 10.3390/ijms241512272.

54. Sharif-Askari, F.S., Sharif-Askari, N.S., Hafezi, S., Mdkhana, B., Alsayed, H.A.H., Ansari, A.W., Mahboub, B., Zakeri, A.M., Temsah, M.-H., Zahir, W., et al. (2022). Interleukin-17, a salivary biomarker for COVID-19 severity. PLOS ONE 17, e0274841. 10.1371/journal.pone.0274841.

55. Ahmed Mostafa, G., Mohamed Ibrahim, H., Al Sayed Shehab, A., Mohamed Magdy, S., AboAbdoun Soliman, N., and Fathy El-Sherif, D. (2022). Up-regulated serum levels of interleukin (IL)-17A and IL-22 in Egyptian pediatric patients with COVID-19 and MIS-C: Relation to the disease outcome. Cytokine 154, 155870. 10.1016/j.cyto.2022.155870.

56. Yuan, S., Jiang, S.C., Zhang, Z.W., Fu, Y.F., Hu, J., and Li, Z.L. (2021). Quantification of Cytokine Storms During Virus Infections. Frontiers in immunology 12, 659419. 10.3389/fimmu.2021.659419.

57. Mahallawi, W.H., Khabour, O.F., Zhang, Q., Makhdoum, H.M., and Suliman, B.A. (2018). MERS-CoV infection in humans is associated with a pro-inflammatory Th1 and Th17 cytokine profile. Cytokine 104, 8–13. 10.1016/j.cyto.2018.01.025.

58. Kelley, L.L., Koury, M.J., Bondurant, M.C., Koury, S.T., Sawyer, S.T., and Wickrema, A. (1993). Survival or death of individual proerythroblasts results from differing erythropoietin sensitivities: a mechanism for controlled rates of erythrocyte production. Blood 82, 2340–2352.

59. Eaves, C.J., and Eaves, A.C. (1978). Erythropoietin (Ep) dose-response curves for three classes of erythroid progenitors in normal human marrow and in patients with polycythemia vera. Blood 52, 1196–1210.

60. Song, X., Dai, D., He, X., Zhu, S., Yao, Y., Gao, H., Wang, J., Qu, F., Qiu, J., Wang, H., et al. (2015). Growth Factor FGF2 Cooperates with Interleukin-17 to Repair Intestinal Epithelial Damage. Immunity 43, 488–501. 10.1016/j.immuni.2015.06.024.

61. Beringer, A., Thiam, N., Molle, J., Bartosch, B., and Miossec, P. (2018). Synergistic effect of interleukin-17 and tumour necrosis factor-alpha on inflammatory response in hepatocytes through interleukin-6-dependent and independent pathways. Clin Exp Immunol 193, 221–233. 10.1111/cei.13140.

62. Doreau, A., Belot, A., Bastid, J., Riche, B., Trescol-Biemont, M.C., Ranchin, B., Fabien, N., Cochat, P., Pouteil-Noble, C., Trolliet, P., et al. (2009). Interleukin 17 acts in synergy with B cell-activating factor to influence B cell biology and the pathophysiology of systemic lupus erythematosus. Nat Immunol 10, 778–785. 10.1038/ni.1741.

63. Hartupee, J., Liu, C., Novotny, M., Li, X., and Hamilton, T. (2007). IL-17 enhances chemokine gene expression through mRNA stabilization. J Immunol 179, 4135–4141. 10.4049/jimmunol.179.6.4135.

64. Pellin, D., Loperfido, M., Baricordi, C., Wolock, S.L., Montepeloso, A., Weinberg, O.K., Biffi, A., Klein, A.M., and Biasco, L. (2019). A comprehensive single cell transcriptional landscape of human hematopoietic progenitors. Nature communications 10, 2395. 10.1038/s41467-019-10291-0.

65. Dahlin, J.S., Hamey, F.K., Pijuan-Sala, B., Shepherd, M., Lau, W.W.Y., Nestorowa, S., Weinreb, C., Wolock, S., Hannah, R., Diamanti, E., et al. (2018). A single-cell hematopoietic landscape resolves 8 lineage trajectories and defects in Kit mutant mice. Blood 131, e1–e11. 10.1182/blood-2017-12-821413.

66. Velten, L., Haas, S.F., Raffel, S., Blaszkiewicz, S., Islam, S., Hennig, B.P., Hirche, C., Lutz, C., Buss, E.C., Nowak, D., et al. (2017). Human haematopoietic stem cell lineage commitment is a continuous process. Nat Cell Biol 19, 271–281. 10.1038/ncb3493.

67. Sathyanarayana, P., Menon, M.P., Bogacheva, O., Bogachev, O., Niss, K., Kapelle, W.S., Houde, E., Fang, J., and Wojchowski, D.M. (2007). Erythropoietin modulation of podocalyxin and a proposed erythroblast niche. Blood 110, 509–518. blood-2006-11-056465 [pii] 10.1182/blood-2006-11-056465.

68. Gillinder, K.R., Tuckey, H., Bell, C.C., Magor, G.W., Huang, S., Ilsley, M.D., and Perkins, A.C. (2017). Direct targets of pSTAT5 signalling in erythropoiesis. PLoS One 12, e0180922. 10.1371/journal.pone.0180922.

69. Kuhrt, D., and Wojchowski, D.M. (2015). Emerging EPO and EPO receptor regulators and signal transducers. Blood 125, 3536–3541. 10.1182/blood-2014-11-575357.

70. Monin, L., and Gaffen, S.L. (2018). Interleukin 17 Family Cytokines: Signaling Mechanisms, Biological Activities, and Therapeutic Implications. Cold Spring Harb Perspect Biol 10. 10.1101/cshperspect.a028522.

71. Eastman, A.E., Chen, X., Hu, X., Hartman, A.A., Pearlman Morales, A.M., Yang, C., Lu, J., Kueh, H.Y., and Guo, S. (2020). Resolving Cell Cycle Speed in One Snapshot with a Live-Cell Fluorescent Reporter. Cell reports 31, 107804. 10.1016/j.celrep.2020.107804.

72. Peslak, S.A., Wenger, J., Bemis, J.C., Kingsley, P.D., Frame, J.M., Koniski, A.D., Chen, Y., Williams, J.P., McGrath, K.E., Dertinger, S.D., and Palis, J. (2011). Sublethal radiation injury uncovers a functional transition during erythroid maturation. Exp Hematol 39, 434–445. S0301-472X(11)00019-1 [pii] 10.1016/j.exphem.2011.01.010.

73. Maira, D., Duca, L., Busti, F., Consonni, D., Salvatici, M., Vianello, A., Milani, A., Guzzardella, A., Di Pierro, E., Aliberti, S., et al. (2022). The role of hypoxia and inflammation in the regulation of iron metabolism and erythropoiesis in COVID-19: The IRONCOVID study. Am J Hematol 97, 1404–1412. 10.1002/ajh.26679.

74. Somers, V.K., Kara, T., and Xie, J. (2020). Progressive Hypoxia: A Pivotal Pathophysiologic Mechanism of COVID-19 Pneumonia. Mayo Clin Proc 95, 2339–2342. 10.1016/j.mayocp.2020.09.015.

75. Xie, J., Covassin, N., Fan, Z., Singh, P., Gao, W., Li, G., Kara, T., and Somers, V.K. (2020). Association Between Hypoxemia and Mortality in Patients With COVID-19. Mayo Clin Proc 95, 1138–1147. 10.1016/j.mayocp.2020.04.006.

76. Basan, M., Hui, S., Okano, H., Zhang, Z., Shen, Y., Williamson, J.R., and Hwa, T. (2015). Overflow metabolism in Escherichia coli results from efficient proteome allocation. Nature 528, 99–104. 10.1038/nature15765.

77. Szenk, M., Dill, K.A., and de Graff, A.M.R. (2017). Why Do Fast-Growing Bacteria Enter Overflow Metabolism? Testing the Membrane Real Estate Hypothesis. Cell Syst 5, 95–104. 10.1016/j.cels.2017.06.005.

78. Schwarzenberger, P., Huang, W., Ye, P., Oliver, P., Manuel, M., Zhang, Z., Bagby, G., Nelson, S., and Kolls, J.K. (2000). Requirement of Endogenous Stem Cell Factor and Granulocyte-Colony-Stimulating Factor for IL-17-Mediated Granulopoiesis. The Journal of Immunology 164, 4783–4789. 10.4049/jimmunol.164.9.4783.

79. Tan, W., Liu, B., Barsoum, A., Huang, W., Kolls, J.K., and Schwarzenberger, P. (2013). Requirement of TPO/c-mpl for IL-17A-induced granulopoiesis and megakaryopoiesis. Journal of Leukocyte Biology 94, 1303–1308. 10.1189/jlb.1212639.

## References

1. Schirm, S. & Scholz, M. A biomathematical model of human erythropoiesis and iron metabolism. Sci. Rep. 10, 8602 (2020).

2. Li, H. et al. Rate of Progression through a Continuum of Transit-Amplifying Progenitor Cell States Regulates Blood Cell Production. Dev. Cell 49, 118–129.e7 (2019).

3. Koury, M. J. Tracking erythroid progenitor cells in times of need and times of plenty. Exp. Hematol. 44, 653–663 (2016).

4. Koury, M. J. & Bondurant, M. C. The mechanism of erythropoietin action. Am. J. Kidney Dis. 18, 20–23 (1991).

5. Ebert, B. L. & Bunn, H. F. Regulation of the erythropoietin gene. Blood 94, 1864–1877 (1999).

6. Koury, M. J. & Bondurant, M. C. Erythropoietin retards DNA breakdown and prevents programmed death in erythroid progenitor cells. Science 248, 378–381 (1990).

7. Socolovsky, M., Fallon, A. E., Wang, S., Brugnara, C. & Lodish, H. F. Fetal anemia and apoptosis of red cell progenitors in Stat5a-/-5b-/-mice: a direct role for Stat5 in Bcl-X(L) induction. Cell 98, 181–191 (1999).

8. Dolznig, H. et al. Apoptosis Protection by the Epo Target Bcl-XL Allows Factor-Independent Differentiation of Primary Erythroblasts. Curr. Biol. 12, 1076–1085 (2002).

9. Koury, M. J. & Bondurant, M. C. Maintenance by erythropoietin of viability and maturation of murine erythroid precursor cells. J. Cell. Physiol. 137, 65–74 (1988).

10. Liu, Y. et al. Suppression of Fas-FasL coexpression by erythropoietin mediates erythroblast expansion during the erythropoietic stress response in vivo. Blood 108, 123–133 (2006).

11. Peslak, S. A. et al. EPO-mediated expansion of late-stage erythroid progenitors in the bone marrow initiates recovery from sublethal radiation stress. Blood 120, 2501–2511 (2012).

12. Myers, G. et al. Murine erythroid differentiation kinetics in vivo under normal and anemic stress conditions. Blood Adv 7, 5727–5732 (2023).

13. Ganzoni, A., Hillman, R. S. & Finch, C. A. Maturation of the macroreticulocyte. Br. J. Haematol. 16, 119–135 (1969).

14. Wiczling, P., Ait-Oudhia, S. & Krzyzanski, W. Flow cytometric analysis of reticulocyte maturation after erythropoietin administration in rats. Cytometry A 75, 584–592 (2009).

15. Axelrad, A. A., McLeod, D. L., Shreeve, M. M. & Heath, D. S. Properties of cells that produce erythrocytic colonies in vitro. Hemopoiesis in culture 226 (1974).

16. Porpiglia, E., Hidalgo, D., Koulnis, M., Tzafriri, A. R. & Socolovsky, M. Stat5 signaling specifies basal versus stress erythropoietic responses through distinct binary and graded dynamic modalities. PLoS Biol. 10, e1001383 (2012).

17. Peschle, C. et al. Identification and characterization of three classes of erythroid progenitors in human fetal liver. Blood 58, 565–572 (1981).

18. Gregory, C. J. Erythropoietin sensitivity as a differentiation marker in the hemopoietic system: studies of three erythropoietic colony responses in culture. J. Cell. Physiol. 89, 289–301 (1976).

19. Van Putten, L. M. The life span of red cells in the rat and the mouse as determined by labeling with DFP32 in vivo. Blood 13, 789–794 (1958).

